# Mosaic metabolic ageing: Basal and standard metabolic rate age in opposite directions and independent of environmental quality, sex or lifespan in a passerine

**DOI:** 10.1101/844266

**Authors:** Michael Briga, Simon Verhulst

## Abstract

1. Crucial to our understanding of the ageing process is identifying how traits change with age, which variables alter their ageing process and whether these traits associate with lifespan.
2. We here investigated metabolic ageing in zebra finches. We longitudinally monitored 407 individuals during six years and collected 3213 measurements of two independent mass-adjusted metabolic traits: basal metabolic rate (BMR_m_) at thermoneutral temperatures and standard metabolic rate (SMR_m_), which is the same as BMR_m_ but at ambient temperatures below thermoneutrality.
3. BMR_m_ *decreased* linearly with age, consistent with earlier reports. In contrast, SMR_m_ *increased* linearly with age. To the best of our knowledge, this is the first quantification of SMR_m_ ageing, and thereby of the contrast between SMR_m_ and BMR_m_ ageing.
4. Neither metabolic rate nor metabolic ageing rate were associated with individual lifespan. Moreover, experimental manipulations of environmental quality that decreased BMR_m_ and SMR_m_ and shortened lifespan with 6 months (12%) did not affect the ageing of either metabolic trait. Females lived 2 months (4%) shorter than males, but none of the metabolic traits showed sex-specific differences at any age.
5. Our finding that ageing patterns of metabolic rate vary depending on the ambient temperature illustrates the importance of studying ageing in an ecologically realistic setting.
6. Our results add to the mounting evidence that within an organism ageing is an asynchronous process.

## 1. Introduction

Senescence, the decline in fitness due to declining organismal functioning with age, is common in humans, model organisms and in the wild (Nussey et al. 2013; Belsky et al. 2015; Fontana and Partridge 2015). Ageing is the change in traits with age, irrespective of their association with residual reproductive value; for example, human hair may turn grey with age but this by itself does not affect remaining lifespan. Major challenges in ageing research are to identify how organisms change with age, the factors affecting those changes and in particular how those changes affect fitness and remaining lifespan (Christensen et al. 2010; Nussey et al. 2013; Gaillard and Lemaitre 2017; Partridge and Deelen 2018).

Energy metabolism is an essential resource underlying organismal functioning, reproduction and survival and is therefore a key variable in physiology and ecology, including in the study of ageing (Drent and Daan 1980; Burton et al. 2011; Norin and Metcalfe 2019). For example, consistent individual differences in energy turnover can be a key determinant of behavior, reproduction, survival (Biro and Stamps 2010; Burton et al. 2011). Energy metabolism can be quantified in several ways. A tractable component of energy turnover that is often quantified is basal metabolic rate (BMR), i.e. the minimum energy expenditure of a post-absorptive adult animal measured during the rest phase at thermoneutral temperature (IUPS Thermal Commission, 2001; McNab, 1997). How BMR changes with age within indiviuals has been quantified in a variety of species and environments, including in humans, and all studies consistently reported a decline with age (Elliott et al. 2015). Thus BMR, an individual’s minimum energy expenditure at thermoneutrality, declines with age.

While BMR is measured in the thermoneutral zone, most endotherms spend much of their lives at ambient temperatures that are below thermoneutrality. Standard metabolic rate (SMR), which is the same as BMR, except that Ta is below thermoneutrality, may therefore be a metabolic trait with higher ecological relevance, in the sense that it better reflects metabolic rate in natural conditions. It is often implicitly assumed that BMR and SMR vary in parallel across individuals, but we previously showed that BMR and SMR in the zebra finch *Taeniopygia guttata* correlated poorly both between and within-individuals, despite substantial repeatabilities (Briga and Verhulst 2017). How SMR changes with age in endotherms has to our best knowledge not previously been investigated, but our earlier findings show that we cannot assume that it changes with age in a similar way as BMR.

We monitored the ageing of BMR and SMR in a captive population of a small passerine, the zebra finch. The birds used in this study lived in captivity, which may alter metabolism and possibly its association with age compared with to that of free-living animals (Wiersma and Verhulst 2005; Auer et al. 2016; Briga and Verhulst 2017). Indeed, an essential difference between captive and free-living individuals is that free-living animals generally face foraging costs, which can be a key parameter in ageing (Briga and Verhulst 2015a; Speakman et al. 2015). Foraging costs could drive or reinforce many age associated declines in organismal performance in the wild, although this was rarely studied (reviewed in: Clay et al. 2018). Hence, to broaden the range of environments and increase the ecological relevance of our study, we housed the birds in outdoor aviaries, and permanently exposed half of our population to high foraging costs using a manipulation of flight cost per food reward (Koetsier and Verhulst, 2011). We applied a 2×2 design, independently manipulating foraging costs during development and in adulthood. This manipulation shortened lifespan and decreased body mass, BMR_m_ and SMR_m_ in our experimental population (Briga et al., 2017; Briga and Verhulst, 2017; Briga et al. 2019). In our experiment, individuals experiencing high foraging costs during development and in adulthood (HH group) lived on average six months (12%) shorter than all other groups. Females also lived two months shorter than males (4%). Given these findings, it was of interest to investigate whether lifespan variation between and within experimental treatment groups and sexes was associated with ageing rate. If we assume that factors that shorten lifespan also accelerate ageing, a common assumption in ageing research (Williams 1999), we predict that the shortest living experimental group shows the highest rate of metabolic ageing.

## 2. Material & methods

### 2.1 Experimental setup

Birds were reared in either experimentally small broods (with 2 or 3 chicks) or large broods (between 5 and 8 chicks). These brood sizes are within the range observed in wild (Zann 1996). Growing up in large broods increases costly begging beg more, receive less food and have impaired growth (Briga, 2016; Briga et al., 2017). After nutritional independence and before the start of the foraging cost manipulation, i.e. between 35 days till approximately 120 days, young were housed in larger indoor cages with up to 40 other young of the same sex and two male and two female adults. Once adult, birds were subject to a long-term foraging experiment, (Koetsier and Verhulst 2011). Briefly, birds were housed in eight single sex outdoor aviaries (L×H×W 310×210×150 cm) located in Groningen, the Netherlands (53° 13’ 0” N / 6° 33’ 0” E). Food (tropical seed mixture) water, grit and cuttlebone were provided *ad libitum* and the birds received fortified canary food (“egg food”, by Bogena, Hedel, the Netherlands) in weighed portions, which is consistent with the conditions in other facilities (Griffith et al. 2017). Each aviary contained an approximately equal number of birds and to keep densities within aviaries within a limited range, new birds were added regularly to replace those that died. The first batch was 3-24 months old when the experiment started and birds added later were 3 to 4 months old.

### 2.2 Data collection

Between December 2007 and April 2013 we collected a total of 3213 respirometry measurements on 407 individuals. Of those, 1233 measurements on 386 individuals are BMR measurements (Fig. S1). In the thermoneutral zone an organism does not increase its metabolic rate in order to maintain body temperature and for the zebra finch this zone occurs at ambient temperatures between 32°C and 39°C (Briga and Verhulst, 2017; Calder, 1964). The other 1980 measurements on 372 individuals are of standard metabolic rate (Fig. S1), which we measured in the same way as BMR, except that the ambient temperature was below thermoneutrality, between 5°C and 32°C. The vast majority of these measurements were at temperatures between 23and 29 (Fig. S2). BMR and SMR data were collected over an age range from 0.4 months till 7.2 years (Fig. S2), in the same seasons, i.e. mostly in spring and autumn (Fig. S3) and measurements were randomized across sex and experimental treatments.

Metabolic rate was measured overnight using an open flow respirometer situated in a dark acclimatized room. Metabolic rate measurements started close to sunset (mean = 18:10 h; SD 01:17). Up to sixteen individuals were taken from the aviaries and randomly transferred to one of sixteen 1.5 L metabolic chambers. Neither food nor water was available for birds during the metabolic measurements. Birds were weighed before and after each measurement and we used the mean value to capture mass corrected BMR or SMR. Technical details about the equipment can be found in Bouwhuis et al. (2011). In brief, the air-flow through the metabolic chambers was controlled at 25 l h^−1^ by mass-flow controllers (5850S; Brooks, Rijswijk, the Netherlands) calibrated with a bubble flow meter. Air was dried using a molecular sieve (3 Å; Merck, Darmstadt, Germany) and analyzed by a paramagnetic oxygen analyzer (Servomex Xentra 4100, Crowborough, UK). During measurements each metabolic chamber or reference outdoor air was sampled every 8 min for 60s to stabilize measurement levels (Bouwhuis et al. 2011). In each sampling, we measured O_2_ and CO_2_ concentration and oxygen consumption was calculated using Eq. (6) of Hill (1972). An energy equivalent of 19.7 kJ l^−1^ oxygen consumed was used to calculate energy expenditure in watt (W). Metabolic rate was taken to be the minimum value of a 30-min running average, which included 3–6 measurement per individual. The first measurement hour was excluded to minimize potential effects of handling stress and incomplete mixture of air in the metabolic chamber. Body mass for the metabolic rate measurements was calculated as the average of the before and after measurement values.

### 2.3 Statistical analyses

#### 2.3.1 General approach

All analyses were done using a general linear mixed modeling approach with the function ‘lmer’ of the package ‘lme4’ (Bates et al. 2015) in R version 3.5.3 (R Core Team 2019). All analyses included individual as a random intercept and their age term (see below) nested within individuals as a random slope. The random slope quantifies the variation within individual in ageing and is required for the correct estimation of confidence intervals when investigating within individual changes (Schielzeth and Forstmeier 2009). Such models require considerable sample sizes to accurately estimate fixed and random effects (van de Pol 2012) and our data fulfilled those requirements (Fig. S1). We did not have sufficient power to add other random terms to the model (e.g. nesting individual identity in aviary). In all analyses, we found the model best supported by the data, and hence the ‘significant’ predictors, using Burnham and Anderson’s model selection approach (Burnham and Anderson, 2002; Burnham et al., 2011) based on second order Akaike Information Criterion (AICc) with the function ‘dredge’ of the package ‘MuMIn’ (Barton 2019). In brief, this is a hypothesis-based approach that generates, given a global model, subset models that best fit the data. Model fitting should be considered as a continuum for which alternative models within 4 ΔAICc are plausible and become increasingly equivocal up to 14 ΔAICc, after which they become implausible (Burnham and Anderson, 2002; Burnham et al., 2011). For models within 4 ΔAICc, we report coefficients which are the result of model averaging from the function ‘model.avg’ of the package ‘MuMIn’. Confidence intervals of model parameters were estimated with the Wald approximation in the function ‘confint’. Residuals of all final models were normal distributed and without influential data points or outliers.

#### 2.3.2 Daily and seasonal variation

Data for all traits were collected throughout the year. To avoid confounding age patterns with seasonal effects, we corrected for daily and seasonal variation in metabolic rates. To this end, we first investigated for each trait which variables best characterized the effect of seasonal variation in trait values following Briga and Verhulst (2017). We quantified the effects of daily and seasonal variation with and tested the effects of: (i) daylength, (ii) photoperiod, quantified as a dichotomous variable for increasing vs. decreasing daylength, and (iii) the minimum ambient temperature (MinT). Temperature data were collected at the weather station of Eelde, approximately 7 km from the aviaries (http://www.knmi.nl/klimatologie/), where temperature was recorded 1.5 m above ground, every hour with accuracy of 0.1 °C. Temperature data at the weather station reflect well the climate at the aviaries as shown from aviary temperature data collected over the course of the eight years of observation (Briga, 2016). For BMR_m_, model selection approach identified that the photoperiod was the most important covariate (ΔAICc=−54, Table S1A). The next best fitting covariate was daylength, should be included as it worsened the model fit within the reasonable range of 4 AICc (ΔAICc=+3.1, Table S1A; Burnham et al. 2011). There was no support for the other tested covariates (ΔAICc≥+8.9, Table S1A). For SMR_m_, we found that both photoperiod and MinT were important (ΔAICc=−32 and ΔAICc=−8.7 respectively, Table S1B). The effect of MinT on trait values can last over a range of timescales (van de Pol and Cockburn 2011; van de Pol et al. 2016). We therefore weighed MinT over a time window as we had found earlier to affect lifespan in our study population (Briga and Verhulst, 2015a). This weighing function included for 77% the MinT within 24 hours before measurement and reached 100% within 5 days. A weighted approach provided a slightly better model fit than using MinT of the day before measurement (ΔAICc=−1.5). Overall, our results show that both BMR_m_ and SMR_m_ increased with shorter and colder days and used the variables that best captured this effect to correct for daily and seasonal variation in both metabolic traits.

#### 2.3.3 Metabolic ageing

Population level associations between trait values and age can be composed of two processes: (i) a within individual change in trait value with age and (ii) a between individual change due to selective mortality of individuals with certain trait values. We distinguished the contributions of these two processes using a within subjects centering approach (van de Pol and Verhulst, 2006; van de Pol and Wright, 2009). In this approach the within individual changes are captured in a Δage term, which is the age at measurement mean centered per individual. Models with and without mean-centering age gave similar conclusions (results not shown), but mean-centering age always improved the model fit (BMR_m_: ΔAICc=9.8; SMR_m_: ΔAICc=13.5; for the best fitting models in Table S4 A and B respectively). The between individual change is captured by the term lifespan, mean centered across our population. For censored birds (N=71, see below), lifespan is unknown and hence these received the mean-centered lifespan of zero. These birds thus contribute only to within and not to between individual trait change. In this formulation, selective disappearance occurs when the coefficients differ within and between individuals, which can be statistically tested with the lifespan term in a model with age and lifespan (van de Pol and Verhulst, 2006; van de Pol and Wright, 2009), or more directly with survival analyses, which provides straightforward ways to test whether this disappearance varies between the experimental groups or sexes (see below).

Within individual changes can follow different age trajectories, including no change with age or a change that was linear, quadratic, terminal (i.e. before death) or the combination of linear and terminal. We tested terminal changes by adding a terminal term, coded as a binomial factor for whether or not an individual died within the year following the measurement. Age trajectories can also be more complex following threshold models (Douhard et al. 2017; Briga et al. 2019). We here tested these models, but we lacked the power to include random slopes, to distinguish between one and two threshold models or to identify threshold ages within a biologically relevant window. We therefore limited the description of these models in the supplementary materials to show that threshold models supported the conclusions made by the linear models (Tables S10-S12).

We tested whether (i) sex and (ii) the environmental manipulations affected metabolic ageing by including the interaction between the age terms and sex or experimental manipulations. Sex was always coded as a two-level factor. Effects of the environmental manipulations were tested using (i) three-way interactions (e.g. Δage*development*adult) and (ii) by grouping experimental group in one factor with four levels. Thirdly, because the HH group is the only group that showed reduced lifespan relative to all other groups, we also coded experimental group as a two level factor, i.e. with HH as one level relative to the three other groups (BB, BH, HB). These three approaches to test for an effect of the environmental manipulations gave consistent conclusions. Interactions between sex and the environmental manipulations and higher order interactions between sex and the environmental manipulation age terms were never significant and we do not show them here.

#### 2.3.4 Associations with lifespan

To investigate the association between mass and survival we used Cox Proportional Hazard analyses (CPH) using the function ‘coxme’ of the package ‘coxme’ (Therneau 2019). To avoid pseudo-replication by repeated measurements, we used the first measurement of each individual. These BMR and SMR values were corrected for mass, temporal and seasonal covariates. We then used the residuals of these values for the CPH analyses. In the CPH models we included sex and the four experimental groups as covariates, as these are known to affect lifespan (Briga et al. 2017, Briga et al. 2019). In all these analyses, aviary was included as a random intercept to correct for the joint housing of birds. Of the 407 individuals for which we have a metabolic rate measurement, we monitored 336 till natural death and right censored (i) 48 individuals that were still alive after 8 years of monitoring and (ii) 23 individuals that died by accident or were euthanized for welfare considerations. These 8 years of monitoring are consistent with the study on body mass ageing in Briga et al. (2019). CPH analyses require predictors to be proportional which was the case as indicated by the ‘cox.zph’ function (*X*^2^=2.49, p>0.11).

## 3. Results

### 3.1 Basal metabolic rate

We first investigated the age trajectory of BMR_m_ within individuals. We tested for changes with age that were either linear, quadratic terminal or a combination of linear and terminal. The best fitting shape was a linear decline with age (ΔAICc<−15.7; Fig. 1A; Table S4A). Individuals lost on average 0.0024 W/Yr (−0.0034<95CI<−0.0015). The variation between individuals in within-individual change with age (i.e. random slopes) was small compared to the variation between individuals in BMR_m_ (i.e. random intercept), with respectively a variance explained of 0.7% vs. 26%. Thus, BMR_m_ decreased linearly with age.

**Fig. 1.**
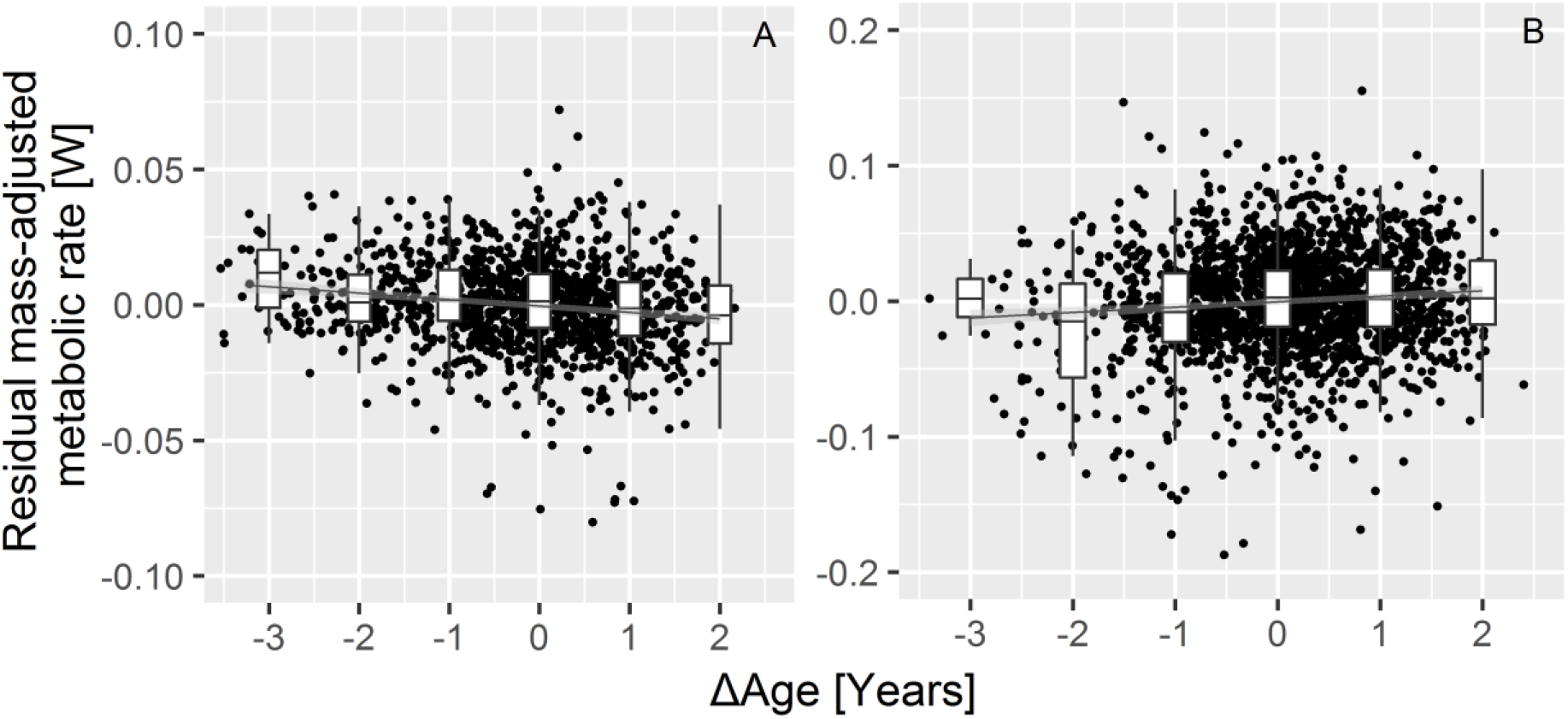
Within individuals (A) BMR_m_ decreased with age, while (B) SMR_m_ increased. Values are residuals corrected for mass, seasonal variation, experimental groups, sex and for SMR_m_ also the ambient temperature at measurement. Boxplots show medians, quartiles and 95% CI per year. Horizontal lines illustrate model fit. Note the Y-axes on different scales between (A) and (B).

We then investigated whether the environmental manipulations that shortened lifespan, also accelerated BMR_m_ decline. High foraging costs during adulthood, but not during development decreased BMR_m_ (development: ΔAICc=+9.7; adulthood: ΔAICc=−12.6; Table S2). There was no evidence for different age trajectories (Table S4A) or for a difference in the magnitude of the linear decline between experimental groups (Fig. 3A-D; Δage * manipulation: ΔAICc>+13.8; terminal year * manipulation: ΔAICc>+21.4; Table S5). Testing the HH group against all other experimental groups gave supported this conclusion (ΔAICc>+13.5). Females lived on average two months shorter than males (4%; Briga et al., 2019). However, sexes did not differ in their BMR_m_ (ΔAICc=+4.7; Table S2), in their BMR_m_ response to the environmental manipulations (ΔAICc>+11.6; Table S2) or in their BMR_m_ ageing (Fig. 3 E&F; ΔAICc>+19.2; Table S7). Thus, BMR_m_ decreased linearly with age and this decrease was independent of factors that affected lifespan.

We then investigated the association between BMR_m_ and lifespan. In the best fitting general linear model, the gradual decline in BMR_m_ with age was indistinguishable between and within individuals (slopes: −0.0018 W/Yr vs. −0.0024 W/Yr respectively), indicating no selective disappearance in any of the experimental groups (Table S5). This is confirmed with CPH models showing neither a linear nor quadratic association with lifespan (e.g. when individuals with intermediate BMR_m_ live longest; Fig. 2; AICc>+1.3; Table S9A). Thus, there was no association between BMR_m_ and lifespan.

**Fig. 2.**
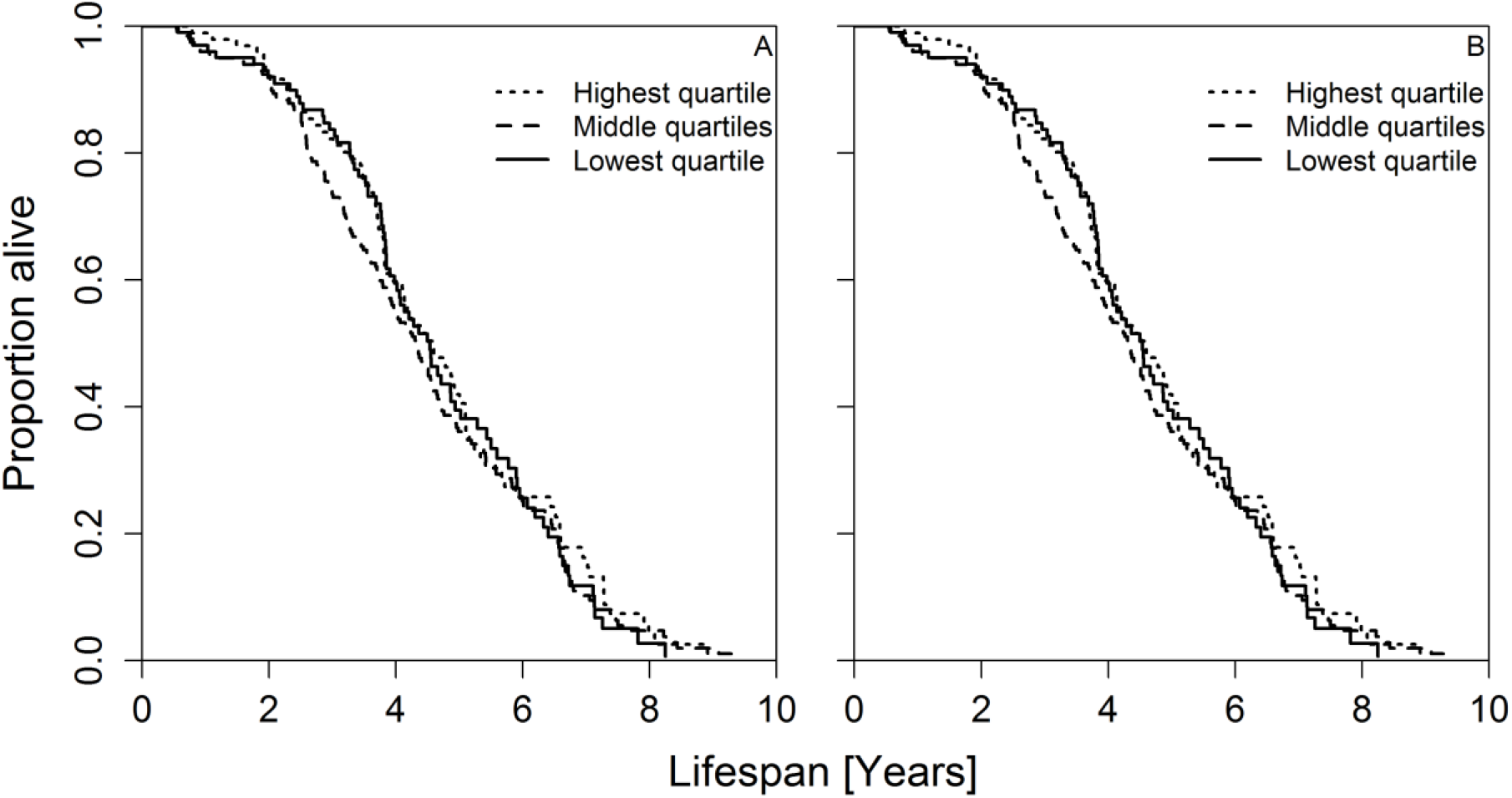
BMR_m_ (A) or SMR_m_ (B) were not associated with lifespan. To avoid pseudo-replication by repeated measurements, we used the first measurement of each individual. BMR_m_ or SMR_m_ values are residuals corrected for mass, seasonal variation, ambient temperature at measurement (SMR_m_), experimental groups and sex, as in the survival analyses in Table S9.

### 3.2 Standard metabolic rate

We first investigated the shape of the age trajectory, testing for linear, quadratic and/or terminal changes. In contrast to BMR_m_, which *decreased* linearly with age, SMR_m_ *increased* linearly with age (ΔAICc<−4.0; Fig. 1B; Table S4B). Individuals gained on average 0.0042 W/Yr (0.0019<95CI<−0.0063). This coefficient seemed to increase with lower ambient temperature Ta at measurement, but this interaction did not improve the model (AICc=+11). The variation between individuals in within-individual change with age was small compared to the variation between individuals in SMR_m_ with respectively a variance explained of 1.5% vs. 20%.

We then investigated whether the manipulations that affected lifespan also affected SMR_m_ ageing. High foraging costs during adulthood, but not during development decreased SMR_m_ (development: ΔAICc=+10.5; adulthood: ΔAICc=−67.6; Table S3). There was no evidence for different age trajectories (Table S4B) or in the magnitude of the increase between experimental groups (Fig. 3 G-J; Δage * manipulation: ΔAICc>+8.1; terminal year * manipulation: ΔAICc>+16.1; Table S6). Testing the HH group against all other experimental groups supported this conclusion (ΔAICc>+11.8). Just as for BMR_m_, sexes did not differ in their SMR_m_ (ΔAICc=+11.8; Table S3), in their SMR_m_ response to the environmental manipulations (ΔAICc>+21.9; Table S3), in their age trajectories or in their rate of SMR_m_ ageing (ΔAICc>+22.1; Table S8). Thus, SMR_m_ increased linearly with age and this increase was independent of factors that affected lifespan.

**Fig. 3.**
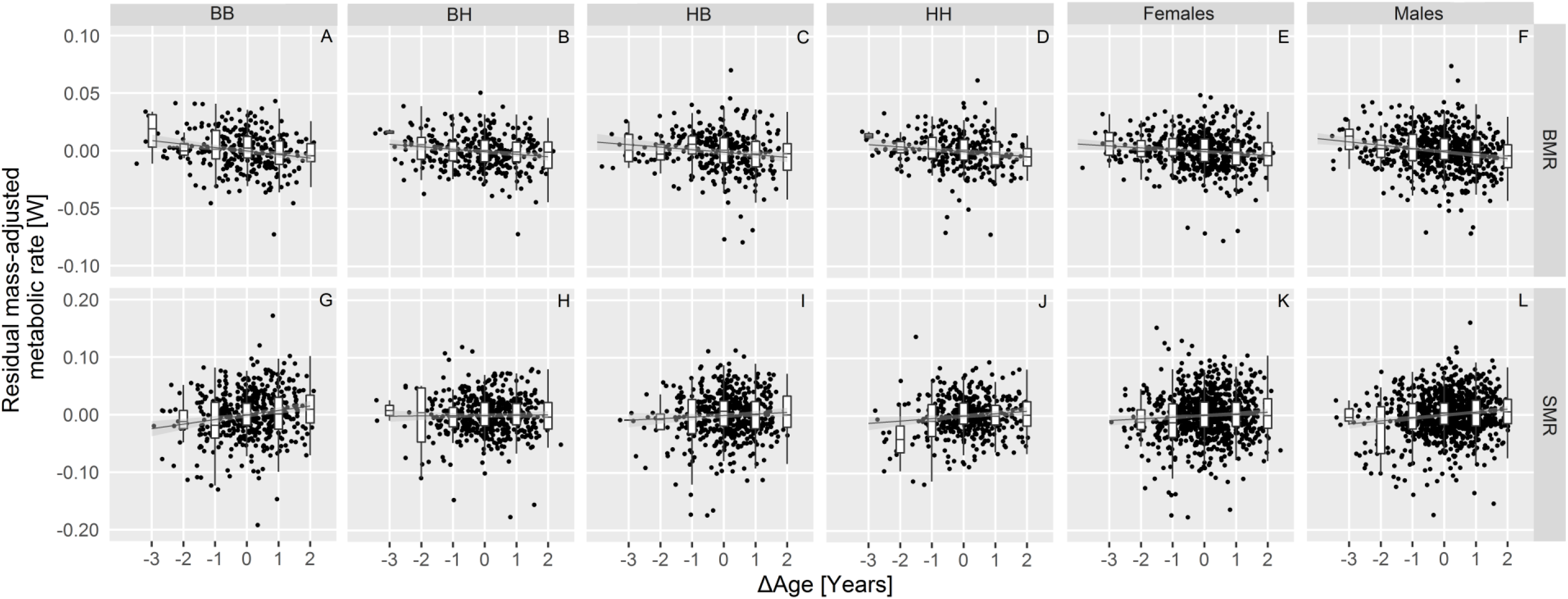
Within-individual changes in BMR_m_ and SMR_m_ are independent of environmental quality (BMR_m_: A-D; SMR_m_: G-J) and of sex (BMR_m_: E, F; SMR_m_: K, L). Experimental group combinations are abbreviated, in chronological order, such that the first letter stands for developmental environment (B for benign or small broods, H for harsh or large broods) and the second letter for adult environment (B for benign or low foraging costs, H for harsh or high foraging costs). BMR_m_ or SMR_m_ values are residuals corrected for mass, seasonal variation, ambient temperature at measurement (SMR_m_), sex (A-D; G-J), experimental groups (E, F, K, L). Boxplots show medians, quartiles and 95% CI per year. Horizontal lines are group specific model fits (± 95%CI).

Lastly, we investigated the association between SMR_m_ and individual variation in lifespan. In a GLM, the gradual increase in SMR_m_ with age differed somewhat between and within individuals (0.0022 W/Yr vs. 0.0042 W/Yr respectively; Table S6), which suggests there could be some selective disappearance with respect to SMR_m_. We found however no statistical support for selective disappearance in a linear model with age and lifespan (ΔAICc=+12.1) or for linear or quadratic association between SMR_m_ and lifespan in a CPH analyses (Fig. 2; ΔAICc>+1.7; Table S9B). Thus, we found no evidence for an association between SMR_m_ and lifespan.

## 4. Discussion

Metabolic rate is a key physiological trait in ecology, life history and the biology of ageing. As such it is common to quantify the minimum energy expenditure at thermoneutral temperatures or BMR_m_. We found that BMR_m_ *decreased* with age, a result consistent with that in many other endotherms, including in zebra finches (Moe et al. 2009; Rønning et al. 2014; Elliott et al. 2015). However, in contrast to BMR_m_, SMR_m_ *increased* with age, at a rate twice the absolute value of BMR_m_ decline. To the best of our knowledge, this is a new result and the discrepancy in ageing trajectory between these two energetic traits is important because it indicates that these change with age independently. Indeed, the correlations between BMR_m_ and SMR_m_ within individual were low (0.04<r<0.22; Briga and Verhulst, 2017) and these traits’ independent ageing will have contributed to this correlation being low. An experimental manipulation of increased foraging costs that decreased lifespan also decreased BMR_m_ and SMR_m_, a result consistent with that of earlier studies in birds and laboratory rodents (Wiersma and Verhulst 2005; Vaanholt et al. 2007; Schubert et al. 2008, 2009). However metabolic rate or metabolic ageing were not associated with lifespan in our study. We here discuss these results in the light of other studies on metabolism and ageing.

### 4.1 Metabolic ageing

The mechanism underlying the change in BMR_m_ with age is likely reductions in the size of metabolic expensive organs, such as heart, liver, kidneys and muscle, as was shown for humans and laboratory rodents (Roberts and Rosenberg 2006). There is also evidence for a small age associated decline in metabolism per unit of tissue (Roberts and Rosenberg 2006) and for reductions in body temperature (Florez-Duquet and McDonald 1998; Weinert 2010; Blatteis 2012). The difference in age trajectory between BMR_m_ and SMR_m_ indicates that there is also ageing of thermoregulation. We suggest two explanations for the increase in SMR_m_ with age. One possibility is that birds need more energy to maintain body temperature as they age, e.g. due to poorer insulation or reduced metabolic efficiency. Secondly, birds in our study have lower body temperatures after nights at sub-thermoneutral temperatures (Briga & Verhulst, 2017), and birds might have developed lower tolerance for low body temperature as they age. This might occur because the ability to warm up decreases with age, at least in laboratory rodents (Florez-Duquet and McDonald 1998). Note however that birds differ from laboratory rodents in that thermoregulatory responses are regulated by skeletal muscles while mammals use brown adipose tissue (Dawson and O’Connor 1996; Mezentseva et al. 2008). Thus, we propose that the mechanisms underlying the different age trajectories between BMR_m_ and SMR_m_ reflect changes in insulation and efficiency of heat production, which is key in the light of the energetic and thermoregulatory challenges associated with winter, predation or with high workload (Brodin 2007; Newton 2007; Speakman and Krol 2010; Nord and Nilsson 2018; Andreassen et al. 2019).

### 4.2 Mosaic ageing and the association with lifespan

Williams (1957) and Maynard-Smith (1962), suggested that traits would evolve to age in synchrony, because “*natural selection will always be in greatest opposition to the decline of the most senescence-prone system*” (Williams 1957). Our findings run counter to this prediction because mass, BMR_m_ and SMR_m_ showed different age trajectories. Moreover, we previously showed mass (Briga et al. 2019) and bill colour (Simons et al. 2012; Simons et al. 2016) in the same population to follow ageing trajectories that differed from both metabolic traits. The age trajectory of mass was quadratic, but this varied with foraging costs in females, but not in males, and bill color, a sexual signal, remained constant through adult life until a decline in the terminal year. Our combined results thus show that traits age asynchronously in zebra finches, a phenomenon coined ‘*mosaic ageing*’ (Cevenini et al. 2008; Walker and Herndon 2010) which has also been observed in other systems (Herndon et al. 2002; Bansal et al. 2015; Belsky et al. 2015; Hayward et al. 2015), but remains to be explained by evolutionary theory. The variation between traits in the net fitness costs of changes in trait value (accounting for costs of maintaining trait level) has been proposed as explanation for mosaic ageing, with traits with larger fitness effects predicted to be better ‘canalised’, i.e. change less with age (Boonekamp et al. 2018), but this hypothesis remains to be tested in the context of ageing.

Metabolic rate (BMR, SMR) is considered a key trait in the study of life-history, ecology and evolution (Drent and Daan 1980; Burton et al. 2011), but at the same time studies in endotherms have not revealed consistent relationships between BMR and fitness (components). In humans, high BMR_m_ is associated with increased mortality risk (Ruggiero et al. 2008). In our study, we found no consistent associations between lifespan and either BMR or SMR, and neither was the rate of ageing in these metabolic traits associated with lifespan. On the one hand, this is reason to question whether variation in BMR and SMR between and within individuals can be considered of functional importance. On the other hand, because these associations are correlational, we cannot rule out that experimental manipulations of metabolic rates (e.g. through pharmaceutic treatment) would result in clear fitness effects. Nevertheless, it seems likely that other physiological pathways are more important in mediating, for example, the effect of environmental conditions on lifespan. Some promising results indicate that glucose metabolism, the insulin pathway and the associated glucocorticoid levels are good candidates, although none of these provides a complete picture yet (Jimeno et al. 2017, 2018; Montoya et al. 2018; Regan et al. 2019). Hence other physiological mechanisms such as oxidative stress and the immune system also warrant further study (De Coster et al. 2011; Simons et al. 2014; Speakman et al. 2015).

It is often implicitly assumed that factors changing lifespan will inextricably also alter ageing rate. To what extent and for what kind of traits this is the case however remains to be identified (Bansal et al., 2015; Christensen et al., 2009; Williams, 1999). Indeed, in our study system we found little evidence for such association: the environmental factors affecting lifespan did not affect the ageing of metabolic traits or body mass (Briga et al. 2019), and neither were rates of ageing higher in shorter lived individuals. Apparently, (environmental) factors affecting lifespan can be distinct from factors affecting ageing rate, which was also suggested for genetic factors (Burger and Promislow 2006). Predicting when or whether an environmental variable that alters lifespan will also affect the ageing of some traits is therefore a current challenge.

