## Supplementary figures and tables for "Mosaic metabolic ageing: Basal and standard metabolic rate age in opposite directions and independent of environmental quality, sex or lifespan in a passerine"

### Table of contents

**1. Data distributions across individuals, age and season** **2**

Figs S1, S2, S3

**2. Seasonal variation in metabolic rate** **5**

Table S1

**3. Metabolic rate, sex and environmental quality** **7**

Tables S2, S3

**4. Identifying the best fitting age trajectories** **10**

Table S4

**5. Metabolic ageing is independent of environmental quality** **11**

Tables S5, S6

**6. Metabolic rate and metabolic ageing are independent of sex** **13**

Tables S7, S8

**7. No association between metabolic rate and lifespan** **15**

Table S9

**8. Threshold models support the conclusions of linear models on $BMR_m$ and $SMR_m$** **16**

Tables S10, S11, S12

**Supplementary Information 1: Data distributions across individuals, age and season**

Fig. S1. Distribution of number of birds with their number of measurements for BMR and SMR.

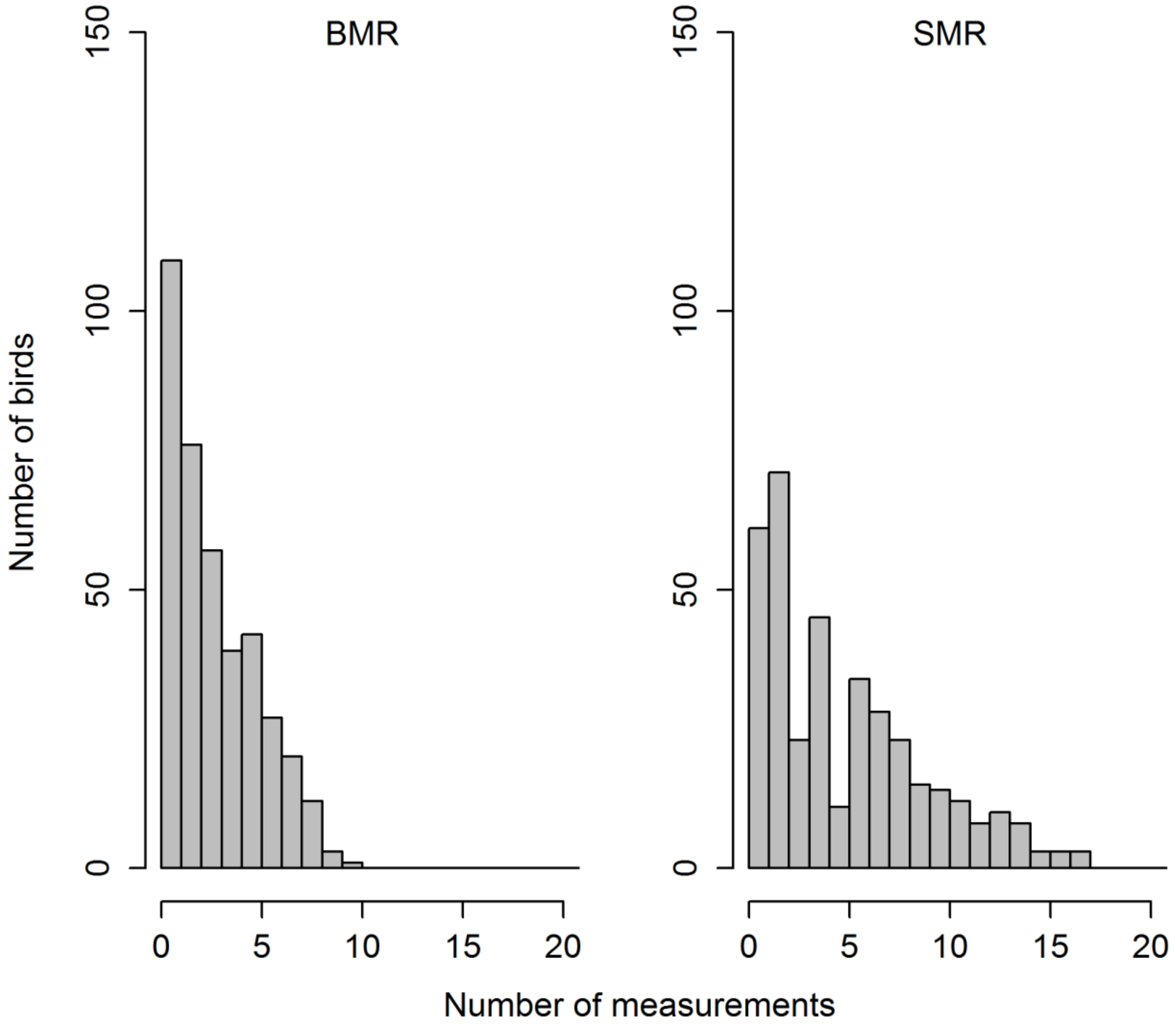

Fig. S2. Distribution of metabolic rate measurements as a function of age, with for BMR ambient temperatures ($T_A$) between 32°C< $T_A$<39°C and for SMR $T_A$<32°C.

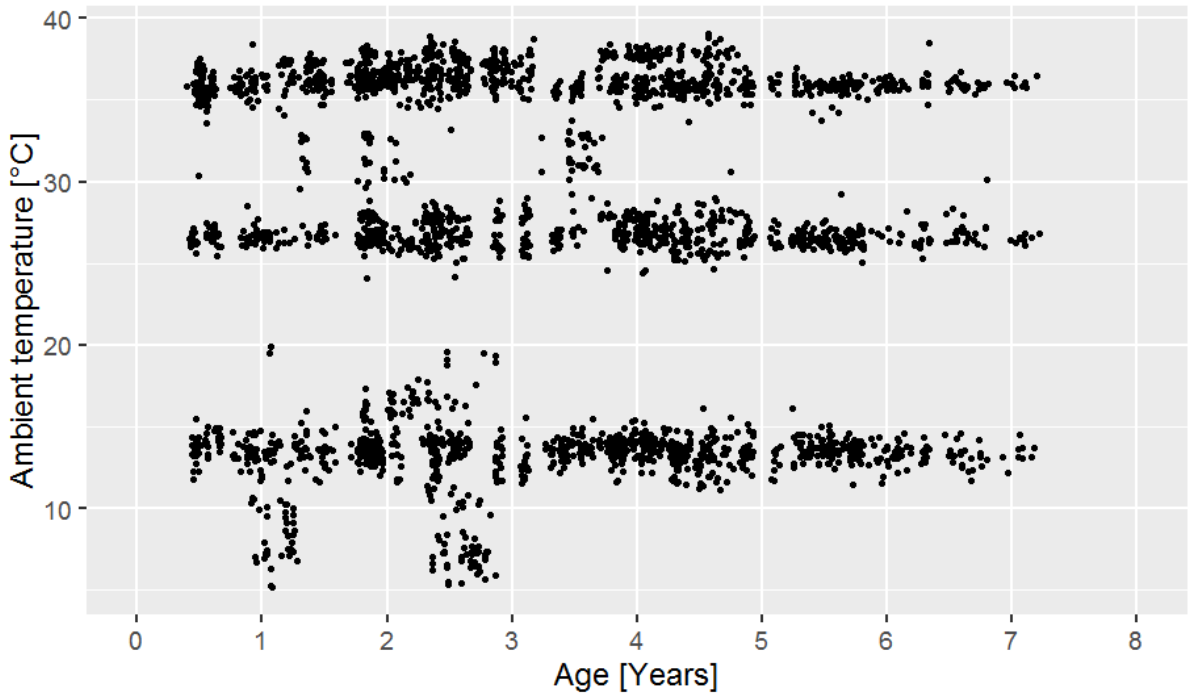

Fig. S3. Monthly distribution of number of measurements for BMR and SMR.

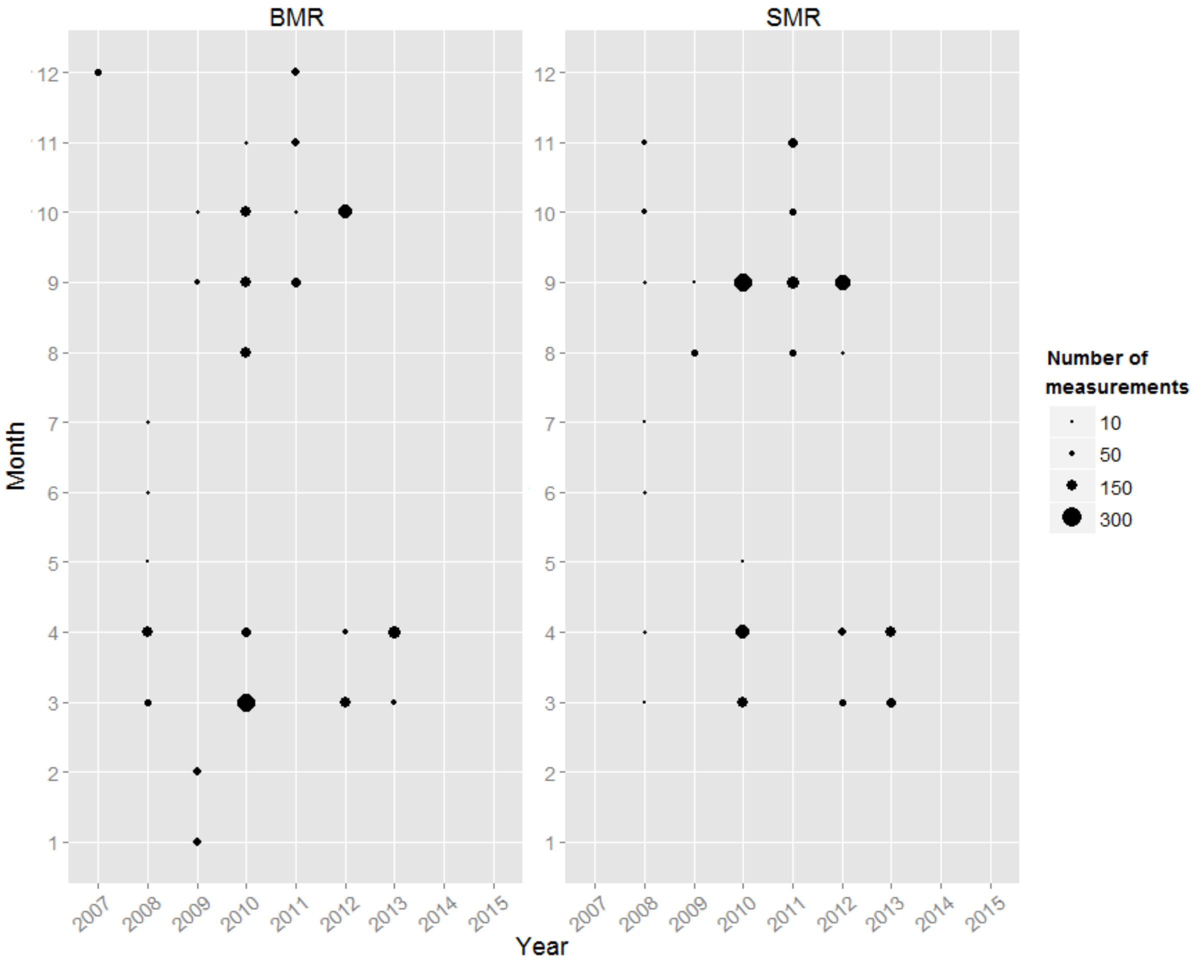

**Supplementary Information 2: Seasonal variation in metabolic rate**

To avoid confounding seasonal variation with age effects, we first investigated which seasonal covariates affected $BMR_m$ and $SMR_m$.

**Basal metabolic rate**

Basal metabolic rate (BMR) was collected from 2008 till 2013 mostly during spring and autumn (Fig. S3). To avoid the confounding effect of mass, we included mass as a covariate in all analyses and hence report mass adjusted BMR ($BMR_m$). $BMR_m$ was higher in spring relative to autumn (photoperiod ΔAICc=-53.6, Table S1A) and increased as days shorten, albeit not significantly (daylength ΔAICc=+3.1). We found no evidence for season specific daylength effects (photoperiod * daylength ΔAICc=+9.8). $BMR_m$ is known to increase in response to colder ambient temperatures on days of or previous to measurement [5]. In our dataset, $BMR_m$ indeed increased with colder ambient MinT (Table S1A), but adding MinT to the model yielded worse model fits both, in addition to or in replacement of daylength and/or season (ΔAICc>+11.9). Thus, $BMR_m$ increased with shorter and colder days, but, in our dataset, seasonal variation in $BMR_m$ was best captured by variables coding for season and, to a lesser extent, daylength. We thus included these covariates in all analyses.

**Standard metabolic rate**

Standard metabolic rate (SMR) was collected just as for BMR, from 2008 till 2013 mostly during spring and autumn (Fig. S3). We here report all mass adjusted SMR ($SMR_m$). Just as for $BMR_m$, $SMR_m$ was higher in spring than in autumn (photoperiod ΔAICc=-32.3, Table S1B) and increased as days shorten (daylength ΔAICc=+1.7) without evidence for season specific daylength effects (photoperiod * daylength ΔAICc=+15.0). SMR also increased on colder days, but, differently from $BMR_m$, the effect of minimum temperature (MinT) on SMR was important (ΔAICc=-8.7). MinT and daylength are correlated and a model with MinT fitted the data better than a model with daylength (ΔAICc=-8.8, Table S1B). Thus, $SMR_m$ increased with shorter and colder days and seasonal variation in $SMR_m$ was best captured by season and MinT. We thus included these covariates in all analyses.

Table S1. Model selection results for time and seasonal effects on mass. $BMR_m$ and $SMR_m$. Abbreviations: Day=Daylength, i.e. the proportion of day between sunrise and sunset; Photo =Photoperiod coded as a dichotomous variable indicating whether daylength increased (0) or decreased (1) relative to the previous day; MinT = Minimum temperature up to 5 days before measurement (see methods); Measm Temp= Ambient temperature of respirometry measurement; df=Number of parameters estimated by the model; AIC=Akaike Information Criterion; ΔAIC=difference in AIC relative to the best fitting model. Models are ordered by increasing AICc. All models include bird identity as random intercept.

| (A) $BMR_m$ | | | Photo | | | | | | |
|---|---|---|---|---|---|---|---|---|---|
| **Model** | **Photo** | **Day** | *** Day** | **MinT** | **Mass** | **df** | **AICc** | **ΔAICc** | **weight** |
| 1 | 0.01 | | | | 0.01 | 5 | -6110.9 | 0 | 0.82 |
| 2 | 0.01 | 0.02 | | | 0.01 | 6 | -6107.8 | 3.1 | 0.17 |
| 3 | 0.02 | 0.03 | -0.02 | | 0.01 | 7 | -6101.1 | 9.8 | 0.01 |
| 4 | 0.01 | | | -0.0003 | 0.01 | 6 | -6099.0 | 11.9 | 0.00 |
| 5 | 0.01 | 0.01 | | -0.0002 | 0.01 | 7 | -6091.8 | 19.1 | 0.00 |
| 6 | 0.02 | 0.02 | -0.02 | -0.0002 | 0.01 | 8 | -6084.5 | 26.5 | 0.00 |
| 7 | | -0.03 | | -0.001 | 0.01 | 6 | -6057.4 | 53.6 | 0.00 |
| 8 | | | | -0.001 | 0.01 | 5 | -6053.4 | 57.5 | 0.00 |
| 9 | | | | | 0.01 | 4 | -6020.8 | 90.1 | 0.00 |
| 10 | | -0.02 | | | 0.01 | 5 | -6017.1 | 93.8 | 0.00 |

| (B) $SMR_m$ | | | Photo | | | Measm | | | | |
|---|---|---|---|---|---|---|---|---|---|---|
| **Model** | **Photo** | **Day** | *** Day** | **MinT** | **Mass** | **Temp** | **df** | **AICc** | **ΔAICc** | **weight** |
| 1 | 0.02 | | | -0.001 | 0.02 | -0.01 | 7 | -6618.1 | 0 | 0.96 |
| 2 | 0.02 | -0.004 | | -0.001 | 0.02 | -0.01 | 8 | -6610.1 | 8.1 | 0.02 |
| 3 | 0.02 | | | | 0.02 | -0.01 | 6 | -6609.5 | 8.7 | 0.01 |
| 4 | 0.03 | 0.04 | | | 0.02 | -0.01 | 7 | -6607.8 | 10.3 | 0.01 |
| 5 | 0.02 | -0.001 | -0.01 | -0.001 | 0.02 | -0.01 | 9 | -6603.1 | 15.0 | 0.00 |
| 6 | 0.04 | 0.05 | -0.02 | | 0.02 | -0.01 | 8 | -6601.2 | 17.0 | 0.00 |
| 7 | | -0.09 | | -0.002 | 0.02 | -0.01 | 7 | -6585.8 | 32.3 | 0.00 |
| 8 | | | | -0.002 | 0.02 | -0.01 | 6 | -6538.6 | 79.5 | 0.00 |
| 9 | | -0.08 | | | 0.02 | -0.01 | 6 | -6497.8 | 120.3 | 0.00 |
| 10 | | | | | 0.02 | -0.01 | 5 | -6470.7 | 147.4 | 0.00 |

### Supplementary information 3: Metabolic rate, sex and environmental quality

#### Basal metabolic rate

BMR was lower in response to both harsh environments but only the harsh adult environment provided a better model fit (development ΔAICc=+1.5; adult ΔAICc=-35.4). There was no evidence for context dependent developmental effects (development * adult ΔAICc=+12.1). Because BMR correlated well with mass (r=0.60), the environmental effects on BMR may be mediated by mass. In order to capture BMR dynamics without the confounding effects of mass, we ran all analyses below with mass as a covariate and refer to this mass adjusted BMR as '$BMR_m$'. Our environmental manipulations affected $BMR_m$ similarly as whole organism BMR, i.e. the harsh adult environment significantly decreased BMR while developmental conditions had no effect (development ΔAICc=+9.7; adult ΔAICc=-12.6; development * adult ΔAICc=+21.3; Table S2). As expected, the effect of the adult environment was more pronounced on whole organism BMR than on $BMR_m$ (-0.016W vs. -0.008W respectively). $BMR_m$ did not differ between the sexes (ΔAICc=+4.7; Table S2), nor did the effect of the environmental manipulations (ΔAICc>+11.6; Table S2), and hence the decrease in on $BMR_m$. Thus, harsh adult but not developmental environments decreased energy consumption at thermoneutrality and this was in part due to lower mass. Birds facing high foraging costs thus also decrease their energy consumption per unit body tissue.

#### Standard metabolic rate

Whole-organism SMR was lower in response to both harsh environments, but only convincingly so for the adult treatment (development ΔAICc=-2.6; adult ΔAICc=-104.9). There was no evidence for context dependent developmental effects (development * adult ΔAICc=+8.5). SMR correlated well with mass (r=0.53, after adjustment for ambient temperature at measurement) and we therefore tested whether these effects remained on SMR adjusted for mass ($SMR_m$). Such a model revealed no effect of developmental manipulation on $SMR_m$ (ΔAICc=+10.5; Table S3), but the effect of the adult manipulation remained (ΔAICc=-67.6; Table S3). As for BMR, the foraging costs effect was more pronounced on whole organism SMR than on $SMR_m$ (-0.045W vs. -0.030W respectively). Interestingly, the foraging cost effect on $SMR_m$ was more than twice that of $BMR_m$ (effect size Cohen d = 0.67 vs. 0.28 respectively). $BMR_m$ did not differ between the sexes (ΔAICc=+11.8; Table S3), nor did the effect of the environmental manipulations (ΔAICc>+21.9; Table S3), and hence the decrease in on $BMR_m$. Thus, both harsh environments decreased energy consumption, but the developmental effect was mediated via mass, while the effect of foraging costs was both, mass dependent and independent.

Table S2. High foraging costs during adulthood decrease $BMR_m$ but there is no effect of the developmental manipulation or sex. Abbreviations: Brood=Brood size manipulation during development; Treat=Foraging treatment manipulation during adulthood; Other abbreviations as in Table S1. All models include bird identity as random intercept.

| $BMR_m$ | Experimental manipulations | | | Sex-specific effects | | | | Covariates | | | Model Fit | | | |
|---|---|---|---|---|---|---|---|---|---|---|---|---|---|---|
| Model | Brood | Treat | Brood * Treat | Sex | Sex * Brood | Sex * Treat | Sex * Brood * Treat | Photo | Day | Mass | df | AICc | ΔAICc | weight |
| 1 | | -0.008 | | | | | | 0.012 | 0.018 | 0.010 | 7 | -6120 | 0.0 | 0.9 |
| 2 | | -0.008 | | -0.005 | | | | 0.012 | 0.018 | 0.010 | 8 | -6116 | 4.7 | 0.1 |
| 3 | -0.003 | -0.008 | | | | | | 0.012 | 0.018 | 0.010 | 8 | -6111 | 9.7 | 0.0 |
| 4 | | -0.012 | | -0.008 | | 0.007 | | 0.012 | 0.018 | 0.010 | 9 | -6109 | 11.6 | 0.0 |
| 5 | | | | | | | | 0.013 | 0.020 | 0.011 | 6 | -6108 | 12.6 | 0.0 |
| 6 | -0.003 | -0.009 | | -0.005 | | | | 0.012 | 0.017 | 0.010 | 9 | -6106 | 14.1 | 0.0 |
| 7 | | | | -0.004 | | | | 0.013 | 0.020 | 0.011 | 7 | -6102 | 18.8 | 0.0 |
| 8 | -0.003 | -0.012 | | -0.008 | | 0.007 | | 0.012 | 0.017 | 0.010 | 10 | -6100 | 20.4 | 0.0 |
| 9 | -0.002 | -0.008 | -0.001 | | | | | 0.012 | 0.018 | 0.010 | 9 | -6099 | 21.3 | 0.0 |
| 10 | -0.002 | | | | | | | 0.013 | 0.020 | 0.011 | 7 | -6097 | 23.5 | 0.0 |
| 11 | -0.004 | -0.009 | | -0.005 | 0.001 | | | 0.012 | 0.017 | 0.010 | 10 | -6095 | 25.7 | 0.0 |
| 12 | -0.003 | -0.008 | -0.001 | -0.005 | | | | 0.012 | 0.017 | 0.010 | 10 | -6095 | 25.8 | 0.0 |
| 13 | -0.002 | | | -0.004 | | | | 0.013 | 0.019 | 0.011 | 8 | -6091 | 29.5 | 0.0 |
| 14 | -0.004 | -0.012 | | -0.009 | 0.001 | 0.007 | | 0.012 | 0.017 | 0.010 | 11 | -6088 | 32.0 | 0.0 |
| 15 | -0.003 | -0.012 | -0.001 | -0.008 | | 0.007 | | 0.012 | 0.017 | 0.010 | 11 | -6088 | 32.2 | 0.0 |
| 16 | -0.003 | -0.008 | -0.001 | -0.005 | 0.001 | | | 0.012 | 0.018 | 0.010 | 11 | -6083 | 37.4 | 0.0 |
| 17 | -0.003 | | | -0.004 | 0.000 | | | 0.013 | 0.019 | 0.011 | 9 | -6079 | 41.2 | 0.0 |
| 18 | -0.004 | -0.012 | -0.001 | -0.009 | 0.001 | 0.007 | | 0.012 | 0.017 | 0.010 | 12 | -6077 | 43.7 | 0.0 |
| 19 | -0.005 | -0.013 | 0.002 | -0.010 | 0.004 | 0.010 | -0.006 | 0.012 | 0.017 | 0.010 | 13 | -6067 | 53.2 | 0.0 |

Table S3. High foraging costs during adulthood decrease $SMR_m$ but there is no effect of the developmental manipulation or sex. Abbreviations: Brood=Brood size manipulation during development; Treat=Foraging treatment manipulation during adulthood; Other abbreviations as in Table S1.

| $SMR_m$ | Experimental manipulations | | | Sex-specific effects | | | | Covariates | | | | Model Fit | | | |
|---|---|---|---|---|---|---|---|---|---|---|---|---|---|---|---|
| Model | Brood | Treat | Brood * Treat | Sex | Sex * Brood | Sex * Treat | Sex * Brood * Treat | Photo | Day | Measm Temp | Mass | df | AICc | ΔAICc | weight |
| 1 | | -0.030 | | | | | | 0.022 | -0.001 | -0.013 | 0.020 | 8 | -6686 | 0.0 | 1.0 |
| 2 | -0.003 | -0.030 | | | | | | 0.022 | -0.001 | -0.013 | 0.020 | 9 | -6675 | 10.5 | 0.0 |
| 3 | | -0.030 | | -0.001 | | | | 0.022 | -0.001 | -0.013 | 0.020 | 9 | -6674 | 11.8 | 0.0 |
| 4 | 0.000 | -0.027 | -0.007 | | | | | 0.022 | -0.001 | -0.013 | 0.020 | 10 | -6666 | 19.5 | 0.0 |
| 5 | | -0.031 | | -0.002 | | 0.003 | | 0.022 | -0.001 | -0.013 | 0.020 | 10 | -6664 | 21.9 | 0.0 |
| 6 | -0.004 | -0.030 | | -0.001 | | | | 0.022 | -0.001 | -0.013 | 0.020 | 10 | -6664 | 22.2 | 0.0 |
| 7 | 0.000 | -0.027 | -0.007 | -0.001 | | | | 0.022 | -0.001 | -0.013 | 0.020 | 11 | -6654 | 31.3 | 0.0 |
| 8 | -0.001 | -0.030 | | 0.002 | -0.005 | | | 0.022 | -0.001 | -0.013 | 0.020 | 11 | -6654 | 31.9 | 0.0 |
| 9 | -0.004 | -0.032 | | -0.003 | | 0.004 | | 0.022 | -0.001 | -0.013 | 0.020 | 11 | -6654 | 32.3 | 0.0 |
| 10 | 0.002 | -0.027 | -0.007 | 0.002 | -0.005 | | | 0.022 | -0.001 | -0.013 | 0.020 | 12 | -6645 | 41.0 | 0.0 |
| 11 | 0.000 | -0.028 | -0.007 | -0.002 | | 0.003 | | 0.022 | -0.001 | -0.013 | 0.020 | 12 | -6644 | 41.4 | 0.0 |
| 12 | -0.001 | -0.032 | | 0.000 | -0.005 | 0.003 | | 0.022 | -0.001 | -0.013 | 0.020 | 12 | -6644 | 42.0 | 0.0 |
| 13 | 0.002 | -0.028 | -0.007 | 0.000 | -0.005 | 0.003 | | 0.022 | -0.001 | -0.013 | 0.020 | 13 | -6635 | 51.1 | 0.0 |
| 14 | 0.006 | -0.025 | -0.014 | 0.004 | -0.012 | -0.003 | 0.013 | 0.022 | -0.001 | -0.013 | 0.020 | 14 | -6627 | 59.0 | 0.0 |
| 15 | | | | | | | | 0.021 | -0.001 | -0.013 | 0.022 | 7 | -6618 | 67.6 | 0.0 |
| 16 | -0.001 | | | | | | | 0.021 | -0.001 | -0.013 | 0.022 | 8 | -6607 | 79.0 | 0.0 |
| 17 | | | | 0.001 | | | | 0.021 | -0.001 | -0.013 | 0.022 | 8 | -6607 | 79.1 | 0.0 |
| 18 | -0.001 | | | 0.001 | | | | 0.021 | -0.001 | -0.013 | 0.022 | 9 | -6595 | 90.4 | 0.0 |
| 19 | 0.002 | | | 0.004 | -0.007 | | | 0.021 | -0.001 | -0.013 | 0.022 | 10 | -6586 | 99.6 | 0.0 |

**Supplementary information 4: Identifying the best fitting metabolic age trajectories**

Table S4. Linear changes with age are the best fitting age trajectory for (A) $BMR_m$ and (B) $SMR_m$. Experimental group (ExpGroup) combinations are abbreviated, in chronological order, such that the first letter stands for developmental environment (B for benign or small broods, H for harsh or large broods) and the second letter for adult environment (B for benign or low foraging costs, H for harsh or high foraging costs). Other abbreviations as in Table S1. Models per experimental group or sex did not include random slopes, hence the significance of their slopes is tested in Tables S5-S8.

| **(A) $BMR_m$** | **All data** | | | | | | **Per Experimental Group** | | | | | | | | | | **Per Sex** | | | | |
|---|---|---|---|---|---|---|---|---|---|---|---|---|---|---|---|---|---|---|---|---|---|
| | | | | | | | | **BB** | | **BH** | | **HB** | | **HH** | | | | **Females** | | **Males** | |
| **Random slope** | **All age terms *** | | | **Δage** | | | **None** | | | | | | | | | | **None** | | | | |
| **Random Intercept** | **Bird ID** | | | **Bird ID** | | | **BirdID** | | | | | | | | | | **BirdID** | | | | |
| **Age trajectory** | **df** | **AICc** | **ΔAICc** | **df** | **AICc** | **ΔAICc** | **df** | **AICc** | **ΔAICc** | **AICc** | **ΔAICc** | **AICc** | **ΔAICc** | **AICc** | **ΔAICc** | | **df** | **AICc** | **ΔAICc** | **AICc** | **ΔAICc** |
| None | 11 | -6094 | 15.8 | 11 | -6094 | 15.8 | see Table S5 | | | | | | | | | | see Table S6 | | | | |
| **Δage** | **14** | **-6110** | **0.0** | **14** | **-6110** | **0.0** | **9** | **-1430** | ***0*** | -1732 | 0 | ***-1440*** | ***0*** | ***-1317*** | ***0*** | | ***11*** | ***-2935*** | ***0*** | ***-3091*** | ***0*** |
| terminal year | 14 | -6090 | 20.4 | 14 | -6102 | 8.0 | 9 | -1425 | 5.5 | ***-1737*** | ***-5.6*** | -1437 | 3.3 | -1315 | 2.0 | | 11 | ***-2937*** | ***-2.7*** | -3075 | 16.3 |
| Δage + terminal year | 18 | -6094 | 15.7 | 15 | -6098 | 12.1 | 10 | -1418 | 11.8 | -1725 | 6.6 | -1429 | 11.0 | -1305 | 11.8 | | 12 | -2925 | 9.8 | -3079 | 12.4 |
| Δage + $Δage^2$ | 18 | -6091 | 19.3 | 15 | -6095 | 15.0 | 10 | -1416 | 14.5 | -1717 | 15.0 | -1427 | 13.5 | -1302 | 14.5 | | 12 | -2919 | 15.2 | -3077 | 14.2 |
| **Models specifications for *** | | | | | | | | | | | | | | | | | | | | | |
| None | BMR~Mass+Nighttime+IncPhoto+Temp+ExpGroup+Sex+Lifespan+(1\|BirdID) | | | | | | | | | | | | | | | | | | | | |
| Δage | BMR~Mass+Nighttime+IncPhoto+Temp+ExpGroup+Sex+Δage+Lifespan+(Δage\|BirdID) | | | | | | | | | | | | | | | | | | | | |
| terminal year | BMR~Mass+Nighttime+IncPhoto+Temp+ExpGroup+Sex+Termin+Lifespan+(Termin\|BirdID) | | | | | | | | | | | | | | | | | | | | |
| Δage + terminal year | BMR~Mass+Nighttime+IncPhoto+Temp+ExpGroup+Sex+Δage+Termin+Lifespan+((Δage+Termin)\|BirdID) | | | | | | | | | | | | | | | | | | | | |
| Δage + $Δage^2$ | BMR~Mass+Nighttime+IncPhoto+Temp+ExpGroup+Sex+Δage+$Δage^2$+Lifespan+((Δage+$Δage^2$)\|BirdID) | | | | | | | | | | | | | | | | | | | | |

| **(B) $SMR_m$** | **All data** | | | | | | **Per Experimental Group** | | | | | | | | | | **Per Sex** | | | | |
|---|---|---|---|---|---|---|---|---|---|---|---|---|---|---|---|---|---|---|---|---|---|
| | | | | | | | | **BB** | | **BH** | | **HB** | | **HH** | | | | **Female** | | **Male** | |
| **Random slope** | **All age terms *** | | | **Δage** | | | **None** | | | | | | | | | | **None** | | | | |
| **Random Intercept** | **Bird ID** | | | **Bird ID** | | | **BirdID** | | | | | | | | | | **BirdID** | | | | |
| **Age trajectory** | **df** | **AICc** | **ΔAICc** | **df** | **AICc** | **ΔAICc** | **df** | **AICc** | **ΔAICc** | **AICc** | **ΔAICc** | **AICc** | **ΔAICc** | **AICc** | **ΔAICc** | | **df** | **AICc** | **ΔAICc** | **AICc** | **ΔAICc** |
| None | 12 | -6647 | 4.0 | 12 | -6647 | 4.0 | see Table S7 | | | | | | | | | | see Table S8 | | | | |
| ***Δage*** | ***15*** | ***-6651*** | ***0.0*** | ***15*** | ***-6651*** | ***0.0*** | ***10*** | ***-1485*** | ***0*** | ***-1936*** | ***0*** | ***-1662*** | ***0*** | ***-1451*** | ***0*** | | ***12*** | ***-3174*** | ***0*** | ***-3399*** | ***0*** |
| terminal year | 15 | -6636 | 15.4 | 15 | -6640 | 10.8 | 10 | -1464 | 20.69 | -1930 | 5.8 | -1654 | 7.2 | -1439 | 11.2 | | 12 | -3162 | 11.9 | -3377 | 22.0 |
| Δage + terminal year | 19 | -6645 | 6.0 | 16 | -6645 | 6.2 | 11 | -1463 | 21.85 | -1918 | 18.0 | -1645 | 16.6 | -1436 | 14.7 | | 13 | -3159 | 15.1 | -3377 | 21.9 |
| Δage + $Δage^2$ | 19 | -6640 | 10.4 | 16 | -6644 | 7.3 | 11 | -1460 | 24.52 | -1916 | 19.8 | -1643 | 18.2 | -1437 | 14.0 | | 13 | -3155 | 19.2 | -3376 | 23.1 |
| **Models specifications for *** | | | | | | | | | | | | | | | | | | | | | |
| None | SMR~Mass+MinT+IncPhoto+Temp+ExpGroup+Sex+Lifespan+(1\|BirdID) | | | | | | | | | | | | | | | | | | | | |
| Δage | SMR~Mass+MinT+IncPhoto+Temp+ExpGroup+Sex+Δage+Lifespan+(Δage\|BirdID) | | | | | | | | | | | | | | | | | | | | |
| terminal year | SMR~Mass+MinT+IncPhoto+Temp+ExpGroup+Sex+Termin+Lifespan+(Termin\|BirdID) | | | | | | | | | | | | | | | | | | | | |
| Δage + terminal year | SMR~Mass+MinT+IncPhoto+Temp+ExpGroup+Sex+Δage+Termin+Lifespan+((Δage+Termin)\|BirdID) | | | | | | | | | | | | | | | | | | | | |
| Δage + $Δage^2$ | SMR~Mass+Nighttime+IncPhoto+Temp+ExpGroup+Sex+Δage+$Δage^2$+Lifespan+((Δage+$Δage^2$)\|BirdID) | | | | | | | | | | | | | | | | | | | | |

**Supplementary information 5: Metabolic ageing is independent of environmental quality**

Table S5. $BMR_m$ shows no evidence for environment-specific age trajectories or environment-specific changes with age as shown here for (A) Δage or (B) terminal year. Results for environment-specific $Δage^2$ effects are even more non-significant (results not shown).

(A) $BMR_m$: No evidence for environment specific delta age effects

| | Experimental manipulations | | | Age terms and interactions | | | | | Other covariates | | | Model Fit | | | |
|---|---|---|---|---|---|---|---|---|---|---|---|---|---|---|---|
| Model | Brood size | Treat | Brood size * Treat | ΔAge | Brood size * ΔAge | Treat * ΔAge | Brood size * Treat * ΔAge | Lifespan | Photo | Night | Mass | df | AICc | ΔAICc | weight |
| 1 | | -0.008 | | -0.002 | | | | -0.002 | 0.010 | 0.010 | 0.011 | 11 | -6139.6 | 0 | 0.99 |
| 2 | -0.003 | -0.008 | | -0.003 | | | | -0.002 | 0.010 | 0.010 | 0.011 | 12 | -6130.4 | 9.23 | 0.01 |
| 3 | | | | -0.003 | | | | -0.002 | 0.011 | 0.012 | 0.011 | 10 | -6128.5 | 11.13 | 0.00 |
| 4 | | -0.008 | | -0.002 | | -0.0004 | | -0.002 | 0.010 | 0.010 | 0.011 | 12 | -6125.8 | 13.81 | 0.00 |
| 5 | -0.002 | -0.007 | -0.001 | -0.003 | | | | -0.002 | 0.010 | 0.010 | 0.011 | 13 | -6118.6 | 20.99 | 0.00 |
| 6 | -0.002 | | | -0.002 | | | | -0.002 | 0.010 | 0.011 | 0.011 | 11 | -6117.9 | 21.69 | 0.00 |
| 7 | -0.003 | -0.008 | | -0.003 | | -0.0004 | | -0.002 | 0.010 | 0.010 | 0.011 | 13 | -6116.6 | 23.06 | 0.00 |
| 8 | -0.003 | -0.008 | | -0.003 | 0.0004 | | | -0.002 | 0.010 | 0.010 | 0.011 | 13 | -6116.5 | 23.15 | 0.00 |
| 9 | -0.002 | -0.007 | -0.001 | -0.003 | | -0.0004 | | -0.002 | 0.010 | 0.010 | 0.011 | 14 | -6104.8 | 34.82 | 0.00 |
| 10 | -0.002 | -0.007 | -0.001 | -0.002 | 0.0004 | | | -0.002 | 0.010 | 0.010 | 0.011 | 14 | -6104.7 | 34.91 | 0.00 |
| 11 | -0.002 | | | -0.002 | 0.0004 | | | -0.002 | 0.010 | 0.011 | 0.011 | 12 | -6104.0 | 35.66 | 0.00 |
| 12 | -0.003 | -0.008 | | -0.003 | 0.0003 | -0.0004 | | -0.002 | 0.010 | 0.010 | 0.011 | 14 | -6102.6 | 37.02 | 0.00 |
| 13 | -0.002 | -0.007 | -0.001 | -0.002 | 0.0004 | -0.0004 | | -0.002 | 0.010 | 0.010 | 0.011 | 15 | -6090.8 | 48.78 | 0.00 |
| 14 | -0.002 | -0.007 | -0.001 | -0.002 | 0.0011 | 0.0003 | -0.002 | -0.002 | 0.010 | 0.010 | 0.011 | 16 | -6078.6 | 61.04 | 0.00 |

(B) $BMR_m$: No evidence for terminal year effects

| | Experimental manipulations | | | Age terms and interactions | | | | | | Other covariates | | | Model Fit | | | |
|---|---|---|---|---|---|---|---|---|---|---|---|---|---|---|---|---|
| Model | Brood size | Treat | Brood size * Treat | ΔAge | Termin Yr | Brood size * Termin Yr | Treat * Termin Yr | Brood size * Treat *Termin Yr | Lifespan | Photo | Night | Mass | df | AICc | ΔAICc | weight |
| 1 | | -0.008 | | -0.003 | | | | | -0.002 | 0.010 | 0.010 | 0.011 | 11 | -6139.6 | 0 | 0.98 |
| 2 | -0.003 | -0.008 | | -0.003 | | | | | -0.002 | 0.010 | 0.010 | 0.011 | 12 | -6130.4 | 9.23 | 0.01 |
| 3 | | | | -0.003 | | | | | -0.002 | 0.011 | 0.012 | 0.011 | 10 | -6128.5 | 11.13 | 0.00 |
| 4 | | -0.008 | | -0.002 | -0.001 | | | | -0.002 | 0.010 | 0.010 | 0.011 | 12 | -6127.5 | 12.09 | 0.00 |
| 5 | -0.002 | -0.007 | -0.001 | -0.003 | | | | | -0.002 | 0.010 | 0.010 | 0.011 | 13 | -6118.6 | 20.99 | 0.00 |
| 6 | | -0.006 | | -0.002 | 0.001 | | -0.004 | | -0.002 | 0.010 | 0.011 | 0.011 | 13 | -6118.3 | 21.35 | 0.00 |
| 7 | -0.002 | | | -0.003 | | | | | -0.002 | 0.010 | 0.011 | 0.011 | 11 | -6117.9 | 21.69 | 0.00 |
| 8 | -0.003 | -0.008 | | -0.002 | -0.001 | | | | -0.002 | 0.010 | 0.010 | 0.011 | 13 | -6117.6 | 22.02 | 0.00 |
| 9 | | | | -0.003 | -0.001 | | | | -0.002 | 0.011 | 0.012 | 0.011 | 11 | -6116.1 | 23.54 | 0.00 |
| 10 | -0.003 | -0.007 | | -0.002 | 0.001 | | -0.004 | | -0.002 | 0.010 | 0.011 | 0.011 | 14 | -6108.2 | 31.39 | 0.00 |
| 11 | -0.003 | -0.008 | | -0.002 | -0.002 | 0.002 | | | -0.002 | 0.010 | 0.010 | 0.011 | 14 | -6106.7 | 32.95 | 0.00 |
| 12 | -0.002 | -0.007 | -0.001 | -0.002 | -0.001 | | | | -0.002 | 0.010 | 0.010 | 0.011 | 14 | -6106.5 | 33.11 | 0.00 |
| 13 | -0.002 | | | -0.003 | -0.001 | | | | -0.002 | 0.011 | 0.012 | 0.011 | 12 | -6105.5 | 34.12 | 0.00 |
| 14 | -0.002 | -0.006 | -0.001 | -0.002 | 0.001 | | -0.004 | | -0.002 | 0.010 | 0.011 | 0.011 | 15 | -6096.5 | 43.13 | 0.00 |
| 15 | -0.003 | -0.007 | | -0.002 | 0.000 | 0.002 | -0.004 | | -0.002 | 0.010 | 0.011 | 0.011 | 15 | -6096.5 | 43.17 | 0.00 |
| 16 | -0.003 | -0.007 | -0.001 | -0.002 | -0.002 | 0.002 | | | -0.002 | 0.010 | 0.010 | 0.011 | 15 | -6094.9 | 44.72 | 0.00 |
| 17 | -0.003 | | | -0.003 | -0.002 | 0.002 | | | -0.002 | 0.010 | 0.012 | 0.011 | 13 | -6093.9 | 45.73 | 0.00 |
| 18 | -0.003 | -0.006 | -0.001 | -0.002 | 0.000 | 0.002 | -0.004 | | -0.002 | 0.010 | 0.011 | 0.011 | 16 | -6084.7 | 54.91 | 0.00 |
| 19 | -0.003 | -0.006 | -0.002 | -0.002 | 0.000 | 0.001 | -0.005 | 0.002 | -0.002 | 0.010 | 0.011 | 0.011 | 17 | -6074.0 | 65.62 | 0.00 |

Table S6. $SMR_m$ shows no evidence for environment-specific age trajectories or environment-specific changes with age as shown here for (A) Δage or (B) terminal year. Results for environment-specific $\Delta age^2$ effects are even more non-significant (results not shown).

**(A) $SMR_m$: No evidence for environment specific delta age effects**

| | Experimental manipulations | | | Age terms and interactions | | | | | Other covariates | | | | Model Fit | | | |
|---|---|---|---|---|---|---|---|---|---|---|---|---|---|---|---|---|
| Model | Brood size | Treat | Brood size * Treat | ΔAge | Brood size * ΔAge | Treat * ΔAge | Brood size * Treat * ΔAge | Lifespan | Photo | MinT | Measm Temp | Mass | df | AICc | ΔAICc | weight |
| 1 | | -0.030 | | 0.004 | | | | 0.002 | 0.022 | -0.001 | -0.013 | 0.019 | 11 | -6682.4 | 0 | 0.98 |
| 2 | | -0.030 | | 0.007 | | -0.005 | | 0.002 | 0.022 | -0.001 | -0.013 | 0.019 | 12 | -6674.2 | 8.18 | 0.02 |
| 3 | -0.003 | -0.030 | | 0.004 | | | | 0.002 | 0.023 | -0.001 | -0.013 | 0.019 | 12 | -6671.9 | 10.53 | 0.01 |
| 4 | -0.003 | -0.030 | | 0.007 | | -0.005 | | 0.002 | 0.022 | -0.001 | -0.013 | 0.019 | 13 | -6663.7 | 18.71 | 0.00 |
| 5 | 0.000 | -0.027 | -0.007 | 0.004 | | | | 0.002 | 0.023 | -0.001 | -0.013 | 0.019 | 13 | -6662.7 | 19.72 | 0.00 |
| 6 | -0.003 | -0.030 | | 0.005 | -0.001 | | | 0.002 | 0.023 | -0.001 | -0.013 | 0.019 | 13 | -6659.6 | 22.82 | 0.00 |
| 7 | -0.0001 | -0.027 | -0.007 | 0.007 | | -0.005 | | 0.002 | 0.022 | -0.001 | -0.013 | 0.019 | 14 | -6654.5 | 27.90 | 0.00 |
| 8 | -0.003 | -0.030 | | 0.007 | -0.001 | -0.005 | | 0.002 | 0.022 | -0.001 | -0.013 | 0.019 | 14 | -6651.5 | 30.87 | 0.00 |
| 9 | 0.0000 | -0.027 | -0.007 | 0.005 | -0.001 | | | 0.002 | 0.023 | -0.001 | -0.013 | 0.019 | 14 | -6650.4 | 32.00 | 0.00 |
| 10 | 0.0000 | -0.027 | -0.007 | 0.007 | -0.001 | -0.005 | | 0.002 | 0.022 | -0.001 | -0.013 | 0.019 | 15 | -6642.4 | 40.06 | 0.00 |
| 11 | 0.0004 | -0.026 | -0.007 | 0.009 | -0.005 | -0.008 | 0.009 | 0.002 | 0.022 | -0.001 | -0.013 | 0.019 | 16 | -6635.0 | 47.39 | 0.00 |
| 12 | | | | 0.004 | | | | 0.002 | 0.021 | -0.001 | -0.013 | 0.022 | 10 | -6612.5 | 69.89 | 0.00 |
| 13 | -0.001 | | | 0.004 | | | | 0.002 | 0.021 | -0.001 | -0.013 | 0.022 | 11 | -6601.1 | 81.29 | 0.00 |
| 14 | -0.001 | | | 0.004 | -0.001 | | | 0.002 | 0.021 | -0.001 | -0.013 | 0.022 | 12 | -6588.8 | 93.57 | 0.00 |

**(B) $SMR_m$: No evidence for environment specific terminal year effects**

| | Experimental manipulations | | | Age terms and interactions | | | | | | Other covariates | | | | Model Fit | | | |
|---|---|---|---|---|---|---|---|---|---|---|---|---|---|---|---|---|---|
| Model | Brood size | Treat | Brood size * Treat | ΔAge | Termin Yr | Brood size * Termin Yr | Treat * Termin Yr | Brood size * Treat * Termin Yr | Lifespan | Photo | MinT | Measm Temp | Mass | df | AICc | ΔAICc | weight |
| 1 | | -0.030 | | 0.004 | | | | | 0.002 | 0.022 | -0.001 | -0.013 | 0.019 | 11 | -6682.4 | 0 | 0.94 |
| 2 | | -0.030 | | 0.006 | -0.007 | | | | 0.001 | 0.022 | -0.001 | -0.013 | 0.019 | 12 | -6676.6 | 5.84 | 0.05 |
| 3 | -0.003 | -0.030 | | 0.004 | | | | | 0.002 | 0.023 | -0.001 | -0.013 | 0.019 | 12 | -6671.9 | 10.53 | 0.01 |
| 4 | | -0.029 | | 0.006 | -0.005 | | -0.004 | | 0.001 | 0.022 | -0.001 | -0.013 | 0.019 | 13 | -6666.3 | 16.07 | 0.00 |
| 5 | -0.003 | -0.031 | | 0.006 | -0.007 | | | | 0.001 | 0.022 | -0.001 | -0.013 | 0.019 | 13 | -6666.1 | 16.34 | 0.00 |
| 6 | -0.0001 | -0.027 | -0.007 | 0.004 | | | | | 0.002 | 0.023 | -0.001 | -0.013 | 0.019 | 13 | -6662.7 | 19.72 | 0.00 |
| 7 | -0.0003 | -0.028 | -0.006 | 0.006 | -0.007 | | | | 0.001 | 0.022 | -0.001 | -0.013 | 0.019 | 14 | -6656.7 | 25.68 | 0.00 |
| 8 | -0.003 | -0.029 | | 0.006 | -0.005 | | -0.004 | | 0.001 | 0.022 | -0.001 | -0.013 | 0.019 | 14 | -6655.8 | 26.65 | 0.00 |
| 9 | -0.002 | -0.031 | | 0.006 | -0.006 | -0.003 | | | 0.001 | 0.022 | -0.001 | -0.013 | 0.019 | 14 | -6655.6 | 26.83 | 0.00 |
| 10 | -0.0003 | -0.026 | -0.006 | 0.006 | -0.005 | | -0.003 | | 0.001 | 0.022 | -0.001 | -0.013 | 0.019 | 15 | -6646.4 | 36.03 | 0.00 |
| 11 | 0.001 | -0.027 | -0.006 | 0.006 | -0.006 | -0.003 | | | 0.001 | 0.022 | -0.001 | -0.013 | 0.019 | 15 | -6646.3 | 36.16 | 0.00 |
| 12 | -0.002 | -0.029 | | 0.006 | -0.004 | -0.003 | -0.004 | | 0.001 | 0.022 | -0.001 | -0.013 | 0.019 | 15 | -6645.3 | 37.08 | 0.00 |
| 13 | 0.001 | -0.026 | -0.006 | 0.006 | -0.004 | -0.003 | -0.004 | | 0.001 | 0.022 | -0.001 | -0.013 | 0.019 | 16 | -6636.0 | 46.45 | 0.00 |
| 14 | 0.002 | -0.025 | -0.009 | 0.006 | -0.002 | -0.007 | -0.007 | 0.007 | 0.001 | 0.022 | -0.001 | -0.013 | 0.019 | 17 | -6627.0 | 55.38 | 0.00 |
| 15 | | | | 0.004 | | | | | 0.002 | 0.021 | -0.001 | -0.013 | 0.022 | 10 | -6612.5 | 69.89 | 0.00 |
| 16 | | | | 0.005 | -0.006 | | | | 0.002 | 0.021 | -0.001 | -0.013 | 0.021 | 11 | -6604.3 | 78.16 | 0.00 |
| 17 | -0.001 | | | 0.004 | | | | | 0.002 | 0.021 | -0.001 | -0.013 | 0.022 | 11 | -6601.1 | 81.29 | 0.00 |
| 18 | -0.001 | | | 0.005 | -0.006 | | | | 0.002 | 0.021 | -0.001 | -0.013 | 0.021 | 12 | -6592.8 | 89.56 | 0.00 |
| 19 | 0.0001 | | | 0.005 | -0.004 | -0.004 | | | 0.002 | 0.021 | -0.001 | -0.013 | 0.021 | 13 | -6582.6 | 99.84 | 0.00 |

### Supplementary Information 6: Metabolic rate and metabolic ageing are independent of sex

Table S7. $BMR_m$ declines with age shows no statistical support for sex- specific Δage, terminal year effects or selective disappearance. Results for sex-specific $\Delta age^2$ effects are even more non-significant (results not shown). Models are ordered by increasing AICc and for simplicity only the 20 best fitting models are shown.

| $BMR_m$ | Experimental manipulations | | Age terms, sex and interactions | | | | | | | Other covariates | | | Model Fit | | | |
|---|---|---|---|---|---|---|---|---|---|---|---|---|---|---|---|---|
| Model | Brood size | Treat | ΔAge | Termin Yr | Sex | Sex * ΔAge | Sex * Termin Yr | Lifespan | Sex * Lifespan | Photo | Night | Mass | df | AICc | ΔAICc | weight |
| 1 | | -0.008 | -0.002 | | | | | -0.002 | | 0.011 | 0.012 | 0.011 | 11 | -6139.6 | 0 | 0.97 |
| 2 | | -0.008 | -0.002 | | -0.003 | | | -0.002 | | 0.011 | 0.011 | 0.011 | 12 | -6130.8 | 8.83 | 0.01 |
| 3 | -0.003 | -0.008 | -0.002 | | | | | -0.002 | | 0.011 | 0.011 | 0.011 | 12 | -6130.4 | 9.23 | 0.01 |
| 4 | | | -0.003 | | | | | -0.002 | | 0.011 | 0.013 | 0.011 | 10 | -6128.5 | 11.13 | 0.00 |
| 5 | | -0.008 | -0.002 | -0.002 | | | | -0.002 | | 0.011 | 0.012 | 0.011 | 12 | -6127.5 | 12.09 | 0.00 |
| 6 | -0.003 | -0.008 | -0.002 | | -0.003 | | | -0.002 | | 0.011 | 0.011 | 0.011 | 13 | -6121.8 | 17.85 | 0.00 |
| 7 | | -0.008 | -0.001 | | -0.004 | -0.002 | | -0.002 | | 0.011 | 0.012 | 0.011 | 13 | -6120.4 | 19.19 | 0.00 |
| 8 | | -0.008 | -0.002 | -0.002 | -0.003 | | | -0.002 | | 0.011 | 0.012 | 0.011 | 13 | -6118.7 | 20.94 | 0.00 |
| 9 | | | -0.003 | | -0.003 | | | -0.002 | | 0.011 | 0.013 | 0.011 | 11 | -6118.6 | 21.02 | 0.00 |
| 10 | -0.003 | -0.008 | -0.002 | -0.002 | | | | -0.002 | | 0.011 | 0.012 | 0.011 | 13 | -6118.3 | 21.35 | 0.00 |
| 11 | -0.002 | | -0.003 | | | | | -0.002 | | 0.011 | 0.013 | 0.011 | 11 | -6117.9 | 21.69 | 0.00 |
| 12 | | -0.008 | -0.002 | | -0.004 | | | -0.001 | -0.001 | 0.011 | 0.011 | 0.011 | 13 | -6117.8 | 21.83 | 0.00 |
| 13 | | | -0.002 | -0.001 | | | | -0.002 | | 0.011 | 0.014 | 0.011 | 11 | -6116.1 | 23.54 | 0.00 |
| 14 | -0.003 | -0.008 | -0.001 | | -0.004 | -0.002 | | -0.002 | | 0.011 | 0.012 | 0.011 | 14 | -6111.5 | 28.17 | 0.00 |
| 15 | -0.003 | -0.008 | -0.002 | -0.002 | -0.003 | | | -0.002 | | 0.011 | 0.011 | 0.011 | 14 | -6109.6 | 29.99 | 0.00 |
| 16 | | -0.008 | -0.001 | -0.002 | -0.004 | -0.002 | | -0.002 | | 0.011 | 0.013 | 0.011 | 14 | -6108.6 | 31.02 | 0.00 |
| 17 | -0.003 | -0.008 | -0.002 | | -0.004 | | | -0.001 | -0.001 | 0.011 | 0.011 | 0.011 | 14 | -6108.6 | 31.05 | 0.00 |
| 18 | | | -0.002 | | -0.004 | -0.002 | | -0.002 | | 0.011 | 0.014 | 0.011 | 12 | -6108.5 | 31.10 | 0.00 |
| 19 | | -0.008 | -0.001 | | -0.004 | -0.002 | | -0.001 | -0.001 | 0.011 | 0.012 | 0.011 | 14 | -6108.5 | 31.17 | 0.00 |
| 20 | -0.003 | | -0.003 | | -0.003 | | | -0.002 | | 0.011 | 0.013 | 0.011 | 12 | -6108.2 | 31.46 | 0.00 |

Table S8. $SMR_m$ declines with age shows no statistical support for sex- specific Δage, terminal year effects or selective disappearance. Results for sex-specific $\Delta age^2$ effects are even more non-significant (results not shown). Models are ordered by increasing AICc and for simplicity only the 20 best fitting models are shown.

| **$SMR_m$** | **Experimental manipulations** | | **Age terms, sex and interactions** | | | | | | | **Other covariates** | | | | **Model Fit** | | | |
|---|---|---|---|---|---|---|---|---|---|---|---|---|---|---|---|---|---|
| **Model** | **Brood size** | **Treat** | **ΔAge** | **TerminYr** | **Sex** | **Sex * ΔAge** | **Sex * TerminYr** | **Lifespan** | **Sex * Lifespan** | **Photo** | **MinT** | **Measm Temp** | **Mass** | **df** | **AICc** | **ΔAICc** | **weight** |
| 1 | | -0.030 | 0.004 | | | | | 0.002 | | 0.022 | -0.001 | -0.013 | 0.019 | 11 | -6682.4 | 0 | 0.94 |
| 2 | | -0.030 | 0.006 | -0.007 | | | | 0.001 | | 0.022 | -0.001 | -0.013 | 0.019 | 12 | -6676.6 | 5.84 | 0.05 |
| 3 | -0.003 | -0.030 | 0.004 | | | | | 0.002 | | 0.023 | -0.001 | -0.013 | 0.019 | 12 | -6671.9 | 10.53 | 0.01 |
| 4 | | -0.030 | 0.004 | | -0.002 | | | 0.002 | | 0.022 | -0.001 | -0.013 | 0.019 | 12 | -6671.0 | 11.46 | 0.00 |
| 5 | -0.003 | -0.031 | 0.006 | -0.007 | | | | 0.001 | | 0.022 | -0.001 | -0.013 | 0.019 | 13 | -6666.1 | 16.34 | 0.00 |
| 6 | | -0.030 | 0.006 | -0.007 | -0.002 | | | 0.001 | | 0.022 | -0.001 | -0.013 | 0.019 | 13 | -6665.2 | 17.22 | 0.00 |
| 7 | -0.003 | -0.030 | 0.004 | | -0.002 | | | 0.002 | | 0.023 | -0.001 | -0.013 | 0.019 | 13 | -6660.5 | 21.95 | 0.00 |
| 8 | | -0.030 | 0.003 | | -0.002 | 0.003 | | 0.002 | | 0.022 | -0.001 | -0.013 | 0.019 | 13 | -6660.3 | 22.14 | 0.00 |
| 9 | | -0.030 | 0.004 | | -0.002 | | | 0.003 | -0.001 | 0.022 | -0.001 | -0.013 | 0.019 | 13 | -6658.3 | 24.14 | 0.00 |
| 10 | | -0.030 | 0.006 | -0.011 | -0.005 | | 0.008 | 0.002 | | 0.022 | -0.001 | -0.013 | 0.019 | 14 | -6657.0 | 25.41 | 0.00 |
| 11 | -0.004 | -0.031 | 0.006 | -0.007 | -0.002 | | | 0.001 | | 0.022 | -0.001 | -0.013 | 0.019 | 14 | -6654.7 | 27.66 | 0.00 |
| 12 | | -0.030 | 0.005 | -0.007 | -0.002 | 0.002 | | 0.002 | | 0.022 | -0.001 | -0.013 | 0.019 | 14 | -6654.0 | 28.40 | 0.00 |
| 13 | | -0.030 | 0.006 | -0.007 | -0.002 | | | 0.002 | -0.001 | 0.022 | -0.001 | -0.013 | 0.019 | 14 | -6652.5 | 29.92 | 0.00 |
| 14 | -0.003 | -0.030 | 0.003 | | -0.002 | 0.003 | | 0.002 | | 0.023 | -0.001 | -0.013 | 0.019 | 14 | -6649.8 | 32.65 | 0.00 |
| 15 | -0.003 | -0.030 | 0.004 | | -0.002 | | | 0.003 | -0.001 | 0.023 | -0.001 | -0.013 | 0.019 | 14 | -6647.7 | 34.67 | 0.00 |
| 16 | | -0.030 | 0.003 | | -0.002 | 0.003 | | 0.003 | -0.001 | 0.022 | -0.001 | -0.013 | 0.019 | 14 | -6647.6 | 34.80 | 0.00 |
| 17 | -0.004 | -0.031 | 0.006 | -0.011 | -0.005 | | 0.008 | 0.002 | | 0.022 | -0.001 | -0.013 | 0.019 | 15 | -6646.6 | 35.85 | 0.00 |
| 18 | | -0.030 | 0.006 | -0.010 | -0.004 | 0.001 | 0.007 | 0.002 | | 0.022 | -0.001 | -0.013 | 0.019 | 15 | -6645.0 | 37.40 | 0.00 |
| 19 | | -0.030 | 0.006 | -0.011 | -0.005 | | 0.008 | 0.001 | 0.000 | 0.022 | -0.001 | -0.013 | 0.019 | 15 | -6644.2 | 38.17 | 0.00 |
| 20 | -0.003 | -0.031 | 0.005 | -0.007 | -0.002 | 0.002 | | 0.001 | | 0.022 | -0.001 | -0.013 | 0.019 | 15 | -6643.6 | 38.86 | 0.00 |

**Supplementary information S7: No association between metabolic rate and lifespan**

Table S9. $BMR_m$ (A) and $SMR_m$ (B) show no association with lifespan and enither was there any evidence for environment-specific or sex-specific association with lifespan. Cox proportional hazard models use residual metabolic rate values which correct for mass, temporal and seasonal covariates as in the best fitting model of table S1. To avoid pseudo-replication by repeated measurements, we used the first measurement of each individual and aviary was included as a random intercept to correct for the joint housing of birds.

| (A) $BMR_m$ | Metabolic terms | | Experiment & Sex Terms | | | | | | Model Fit | | | |
|---|---|---|---|---|---|---|---|---|---|---|---|---|
| | (Residuals) | | | | Exp Group | Exp Group | Sex | Sex | | | | |
| Model | BMR | BMR^2 | ExpGroup | Sex | * BMR | * BMR^2 | * BMR | * BMR^2 | df | AICc | ΔAICc | weight |
| 1 | | | + | + | | | | | 4 | 3202.6 | 0 | 0.42 |
| 2 | -3.40 | | + | + | | | | | 5 | 3203.8 | 1.25 | 0.23 |
| 3 | -0.51 | | + | + | | | + | | 6 | 3205.4 | 2.88 | 0.10 |
| 4 | -3.40 | 1.2 | + | + | | | | | 6 | 3205.9 | 3.31 | 0.08 |
| 5 | 0.25 | | + | + | + | | | | 8 | 3206.8 | 4.27 | 0.05 |
| 6 | -0.51 | 0.5 | + | + | | | + | | 7 | 3207.5 | 4.95 | 0.04 |
| 7 | -0.27 | 266.8 | + | + | | | + | + | 8 | 3207.8 | 5.25 | 0.03 |
| 8 | 0.72 | -76.6 | + | + | + | | | | 9 | 3208.7 | 6.09 | 0.02 |
| 9 | 2.41 | | + | + | + | | + | | 9 | 3208.7 | 6.16 | 0.02 |
| 10 | 2.93 | -77.2 | + | + | + | | + | | 10 | 3210.6 | 8.00 | 0.01 |
| 11 | 1.89 | 173.9 | + | + | + | | + | + | 11 | 3211.3 | 8.75 | 0.01 |
| 12 | 2.09 | -270.3 | + | + | + | + | | | 12 | 3212.9 | 10.32 | 0.00 |
| 13 | 6.93 | -333.9 | + | + | + | + | + | | 13 | 3214.5 | 11.90 | 0.00 |
| 14 | 5.64 | -129.1 | + | + | + | + | + | + | 15 | 3215.8 | 13.19 | 0.00 |
| (B) $SMR_m$ | Metabolic terms | | Experiment & Sex Terms | | | | | | Model Fit | | | |
| | (Residuals) | | | | Exp Group | Exp Group | Sex | Sex | | | | |
| Model | SMR | SMR^2 | ExpGroup | Sex | * SMR | * SMR^2 | * SMR | * SMR^2 | df | AICc | ΔAICc | weight |
| 1 | | | + | + | | | | | 4 | 2959.6 | 0 | 0.42 |
| 2 | -0.77 | | + | + | | | | | 5 | 2961.3 | 1.74 | 0.18 |
| 3 | 4.57 | 29.54 | + | + | + | | | | 9 | 2962.6 | 3.02 | 0.09 |
| 4 | 0.10 | 11.98 | + | + | | | | | 6 | 2962.7 | 3.11 | 0.09 |
| 5 | 0.11 | | + | + | | | + | | 6 | 2963.1 | 3.52 | 0.07 |
| 6 | 1.04 | | + | + | + | | | | 8 | 2963.4 | 3.85 | 0.06 |
| 7 | 5.88 | 27.59 | + | + | + | | + | | 10 | 2964.2 | 4.60 | 0.04 |
| 8 | 2.99 | | + | + | + | | + | | 9 | 2964.6 | 5.00 | 0.03 |
| 9 | 0.53 | 10.57 | + | + | | | + | | 7 | 2964.7 | 5.10 | 0.03 |
| 10 | 6.49 | 48.49 | + | + | + | | + | + | 11 | 2965.6 | 6.00 | 0.02 |
| 11 | 1.62 | 36.92 | + | + | | | + | + | 8 | 2965.9 | 6.30 | 0.02 |
| 12 | 3.73 | 23.08 | + | + | + | + | | | 12 | 2967.6 | 8.01 | 0.01 |
| 13 | 5.22 | 18.44 | + | + | + | + | + | | 13 | 2968.7 | 9.17 | 0.00 |
| 14 | 5.55 | 26.15 | + | + | + | + | + | + | 14 | 2970.9 | 11.30 | 0.00 |

**Supplementary information S8: Threshold models support the conclusions of linear models on $BMR_m$ and $SMR_m$**

Table S10. Description of the threshold models, their model fit and threshold ages for (A) $BMR_m$ and (B) $SMR_m$. Model fits include a penalty of +2AICc per threshold. All differences in threshold ages between experimental groups or sex were within 4 ΔAICc, and hence statistically not distinguishable.

| **$BMR_m$** | **All data** | | **Per Experimental Group** | | | | | **Per Sex** | | |
|---|---|---|---|---|---|---|---|---|---|---|
| **AICc** | **df** | **All** | **df** | **BB** | **BH** | **HB** | **HH** | **df** | **Females** | **Males** |
| 1 threshold | 17 | -6231 | 11 | -1500 | -1807 | -1515 | -1397 | 13 | -3036 | -3198 |
| 2 thresholds | 18 | -6235 | 13 | -1503 | -1808 | -1501 | -1396 | 15 | -3036 | -3202 |
| **ΔAICc** | | | | | | | | | | |
| 1 threshold | 17 | 5 | 10 | 3 | 1 | 0 | 0 | 13 | 0 | 4 |
| 2 thresholds | 18 | 0 | 12 | 0 | 0 | 14 | 1 | 15 | 0 | 0 |
| **Threshold ages** | | | | | | | | | | |
| 1 threshold | NA | 3.1 | NA | 2.4 | 4.3 | 3.2 | 2.2 | NA | 4.8 | 2.3 |
| 2 thresholds, first | NA | 2.8 | NA | 2.8 | 3.5 | 2.8 | 2.7 | NA | 3.5 | 2.3 |
| 2 thresholds, second | NA | 5.0 | NA | 4.8 | 3.9 | 4.8 | 5.7 | NA | 5.1 | 4.8 |
| **Models for all data** | | | | | | | | | | |
| 1 threshold: BMR~Mass+Photo+Day+ExpGroup+Sex+Age1+Age2+Lifespan+(1\|BirdID) | | | | | | | | | | |
| 2 thresholds: BMR~Mass+Photo+Day+ExpGroup+Sex+Age1+Age2+Age3+Lifespan+(1\|BirdID) | | | | | | | | | | |

| **$SMR_m$** | **All data** | | **Per Experimental Group** | | | | | **Per Sex** | | |
|---|---|---|---|---|---|---|---|---|---|---|
| **AICc** | **df** | **All** | **df** | **BB** | **BH** | **HB** | **HH** | **df** | **Females** | **Males** |
| 1 threshold | 18 | -6776 | 12 | -1557 | -2018 | -1739 | -1526 | 14 | -3276 | -3494 |
| 2 thresholds | 19 | -6776 | 14 | -1556 | -2016 | -1736 | -1531 | 16 | -3275 | -3492 |
| **ΔAICc** | | | | | | | | | | |
| 1 threshold | 18 | 0 | 12 | 0 | 0 | 0 | 5 | 14 | 0 | 0 |
| 2 thresholds | 19 | 0 | 14 | 1 | 2 | 3 | 0 | 16 | 1 | 2 |
| **Threshold ages** | | | | | | | | | | |
| 1 threshold | NA | 0.8 | NA | 0.8 | 5.3 | 0.9 | 2.7 | NA | 0.8 | 0.7 |
| 2 thresholds, first | NA | 0.8 | NA | 0.8 | 0.7 | 0.9 | 3.5 | NA | 0.8 | 0.7 |
| 2 thresholds, second | NA | 3.9 | NA | 4.4 | 5.3 | 4.1 | 3.7 | NA | 3.9 | 4.9 |
| **Models for all data** | | | | | | | | | | |
| 1 threshold: SMR~Mass+Photo+MinT+Temp+ExpGroup+Sex+Age1+Age2+Lifespan+(1\|BirdID) | | | | | | | | | | |
| 2 thresholds: SMR~Mass+Photo+MinT+Temp+ExpGroup+Sex+Age1+Age2+Age3+Lifespan+(1\|BirdID) | | | | | | | | | | |

Table S11. Threshold models support the decrease in $BMR_m$ with age without statistical support for sex or environmental manipulations rate of ageing (slopes) for (A) the 1-threshold model and (B) the 2-threshold model, with threshold ages (A) at 3.1 and (B) at 2.8 and 5 years respectively. The coefficients of the models in bold and italic describe a decline in $BMR_m$ with age over most of the age range. For the sake of simplicity, experimental groups and sex were pooled in the 8 level factor group. For (B) we showed only the 15 best fitting models.

| **(A) 1-threshold model** | | | | | | | | | | | | | | | |
|---|---|---|---|---|---|---|---|---|---|---|---|---|---|---|---|
| | **age terms** | | | | **age * group interactions** | | | | **Covariates** | | | **Model fit** | | | |
| | | | | | | **group *** | **group *** | **group *** | | | | | | | |
| **Model** | **age1** | **age2** | **age 3** | **Lifesp** | **group** | **age1** | **age2** | **age 3** | **Photo** | **Day** | **Mass** | **df** | **AICc** | **ΔAICc** | **weight** |
| ***1*** | | | ***NA*** | ***0.002*** | ***+*** | | | ***NA*** | ***0.05*** | ***0.28*** | ***0.01*** | ***14*** | ***-3588*** | ***0*** | ***0.5*** |
| ***2*** | | ***-0.002*** | ***NA*** | ***0.003*** | ***+*** | | | ***NA*** | ***0.05*** | ***0.28*** | ***0.01*** | ***15*** | ***-3587*** | ***1.1*** | ***0.3*** |
| ***3*** | ***0.000*** | | ***NA*** | ***0.002*** | ***+*** | | | ***NA*** | ***0.05*** | ***0.28*** | ***0.01*** | ***15*** | ***-3586*** | ***2.0*** | ***0.2*** |
| ***4*** | ***0.001*** | ***-0.003*** | ***NA*** | ***0.003*** | ***+*** | | | ***NA*** | ***0.05*** | ***0.28*** | ***0.01*** | ***16*** | ***-3585*** | ***3.1*** | ***0.1*** |
| 5 | | -0.005 | NA | 0.003 | + | | + | NA | 0.05 | 0.28 | 0.01 | 22 | -3577 | 11.3 | 0.0 |
| 6 | 0.001 | -0.005 | NA | 0.002 | + | | + | NA | 0.05 | 0.28 | 0.01 | 23 | -3575 | 13.2 | 0.0 |
| 7 | 0.000 | | NA | 0.002 | + | + | | NA | 0.05 | 0.28 | 0.01 | 22 | -3574 | 14.3 | 0.0 |
| 8 | 0.002 | -0.003 | NA | 0.003 | + | + | | NA | 0.05 | 0.29 | 0.01 | 23 | -3573 | 15.3 | 0.0 |
| 9 | 0.004 | -0.006 | NA | 0.002 | + | + | + | NA | 0.05 | 0.29 | 0.01 | 30 | -3562 | 26.3 | 0.0 |
| 10 | | | NA | 0.002 | | | | NA | 0.05 | 0.31 | 0.02 | 7 | -3541 | 47.5 | 0.0 |
| 11 | | -0.002 | NA | 0.003 | | | | NA | 0.05 | 0.31 | 0.02 | 8 | -3539 | 49.1 | 0.0 |
| 12 | 0.000 | | NA | 0.003 | | | | NA | 0.05 | 0.31 | 0.02 | 8 | -3539 | 49.5 | 0.0 |
| 13 | 0.001 | -0.002 | NA | 0.003 | | | | NA | 0.05 | 0.31 | 0.02 | 9 | -3537 | 51.1 | 0.0 |
| **(B) 2-threshold model** | | | | | | | | | | | | | | | |
| ***1*** | | | | ***0.002*** | ***+*** | | | | ***0.05*** | ***0.28*** | ***0.01*** | ***14*** | ***-3588*** | ***0.0*** | ***0.3*** |
| ***2*** | | ***-0.001*** | | ***0.003*** | ***+*** | | | | ***0.05*** | ***0.28*** | ***0.01*** | ***15*** | ***-3587*** | ***1.2*** | ***0.2*** |
| ***3*** | | | ***-0.003*** | ***0.002*** | ***+*** | | | | ***0.05*** | ***0.28*** | ***0.01*** | ***15*** | ***-3586*** | ***1.7*** | ***0.1*** |
| ***4*** | ***0.001*** | | | ***0.002*** | ***+*** | | | | ***0.05*** | ***0.28*** | ***0.01*** | ***15*** | ***-3586*** | ***2.0*** | ***0.1*** |
| ***5*** | ***0.002*** | ***-0.002*** | | ***0.002*** | ***+*** | | | | ***0.05*** | ***0.28*** | ***0.01*** | ***16*** | ***-3585*** | ***2.9*** | ***0.1*** |
| ***6*** | | ***-0.002*** | ***0.001*** | ***0.003*** | ***+*** | | | | ***0.05*** | ***0.28*** | ***0.01*** | ***16*** | ***-3585*** | ***3.2*** | ***0.1*** |
| 7 | ***0.001*** | | ***-0.003*** | ***0.002*** | ***+*** | | | | ***0.05*** | ***0.28*** | ***0.01*** | ***16*** | ***-3584*** | ***3.7*** | ***0.1*** |
| 8 | 0.003 | -0.002 | 0.003 | 0.002 | + | | | | 0.05 | 0.29 | 0.01 | 17 | -3583 | 4.8 | 0.0 |
| 9 | | -0.003 | | 0.003 | + | | + | | 0.05 | 0.28 | 0.01 | 22 | -3576 | 11.9 | 0.0 |
| 10 | | | -0.011 | 0.002 | + | | | + | 0.05 | 0.28 | 0.01 | 22 | -3575 | 13.1 | 0.0 |
| 11 | 0.002 | -0.003 | | 0.002 | + | | + | | 0.05 | 0.28 | 0.01 | 23 | -3575 | 13.6 | 0.0 |
| 12 | | -0.003 | 0.000 | 0.003 | + | | + | | 0.05 | 0.28 | 0.01 | 23 | -3574 | 13.9 | 0.0 |
| 13 | 0.002 | | | 0.002 | + | + | | | 0.05 | 0.29 | 0.01 | 22 | -3573 | 14.6 | 0.0 |
| 14 | | -0.001 | -0.007 | 0.003 | + | | | + | 0.05 | 0.28 | 0.01 | 23 | -3573 | 14.8 | 0.0 |
| 15 | 0.001 | | -0.011 | 0.002 | + | | | + | 0.05 | 0.28 | 0.01 | 23 | -3573 | 15.1 | 0.0 |

Table S12. Threshold models support the increase in $SMR_m$ with age without statistical support for sex or environmental manipulations rate of ageing (slopes) for (A) the 1-threshold model and (B) the 2-threshold model, with threshold ages (A) at 0.8 years and (B) at 0.8 and 3.9 years respectively. The coefficients of the models in bold and italic describe an increase in $SMR_m$ with age over most of the age range. For the sake of simplicity, experimental group and sex were pooled in an 8 level factor. For (B) we showed only the 15 best fitting models.

| **(A) 1-threshold model** | | | | | | | | | | | | | | | | |
|---|---|---|---|---|---|---|---|---|---|---|---|---|---|---|---|---|
| **Model** | **age terms** | | | | **age * group interactions** | | | | **Covariates** | | | | **Model fit** | | | |
| | | | | | | **group *** | **group *** | **group *** | | | **Measm** | | | | | |
| | **age1** | **age2** | **age 3** | **Lifesp** | **group** | **age1** | **age2** | **age 3** | **Photo** | **MinT** | **Temp** | **Mass** | **df** | **AICc** | **ΔAICc** | **weight** |
| 1 | ***-0.076*** | ***0.003*** | ***NA*** | ***0.001*** | ***+*** | | | ***NA*** | ***0.024*** | ***-0.001*** | ***-0.013*** | ***0.019*** | ***17*** | ***-6778*** | ***0.0*** | ***0.9*** |
| 2 | -0.075 | 0.005 | NA | 0.001 | + | | + | NA | 0.02 | 0.00 | -0.01 | 0.02 | 24 | -6772 | 5.9 | 0.0 |
| 3 | -0.146 | 0.003 | NA | 0.001 | + | + | | NA | 0.02 | 0.00 | -0.01 | 0.02 | 24 | -6768 | 10.0 | 0.0 |
| 4 | -0.062 | | NA | 0.003 | + | | | NA | 0.02 | 0.00 | -0.01 | 0.02 | 16 | -6766 | 11.6 | 0.0 |
| 5 | | 0.003 | NA | 0.001 | + | | | NA | 0.02 | 0.00 | -0.01 | 0.02 | 16 | -6765 | 12.5 | 0.0 |
| 6 | -0.164 | 0.006 | NA | 0.001 | + | + | + | NA | 0.02 | 0.00 | -0.01 | 0.02 | 31 | -6762 | 15.8 | 0.0 |
| 7 | | 0.005 | NA | 0.001 | + | | + | NA | 0.02 | 0.00 | -0.01 | 0.02 | 23 | -6760 | 17.9 | 0.0 |
| 8 | | | NA | 0.002 | + | | | NA | 0.02 | 0.00 | -0.01 | 0.02 | 15 | -6758 | 19.6 | 0.0 |
| 9 | -0.129 | | NA | 0.003 | + | + | | NA | 0.02 | 0.00 | -0.01 | 0.02 | 23 | -6756 | 21.5 | 0.0 |
| 10 | -0.076 | 0.004 | NA | 0.001 | | | | NA | 0.02 | 0.00 | -0.01 | 0.02 | 10 | -6707 | 70.9 | 0.0 |
| 11 | | 0.003 | NA | 0.001 | | | | NA | 0.02 | 0.00 | -0.01 | 0.02 | 9 | -6696 | 82.3 | 0.0 |
| 12 | -0.062 | | NA | 0.003 | | | | NA | 0.02 | 0.00 | -0.01 | 0.02 | 9 | -6694 | 84.1 | 0.0 |
| 13 | | | NA | 0.003 | | | | NA | 0.02 | 0.00 | -0.01 | 0.02 | 8 | -6687 | 91.4 | 0.0 |
| **(B) 2-threshold model** | | | | | | | | | | | | | | | | |
| 1 | ***-0.081*** | ***0.004*** | | ***0.001*** | ***+*** | | | | ***0.02*** | ***0.00*** | ***-0.01*** | ***0.02*** | ***17*** | ***-6782*** | ***0.0*** | ***0.7*** |
| 2 | ***-0.080*** | ***0.004*** | ***0.001*** | ***0.001*** | ***+*** | | | | ***0.02*** | ***0.00*** | ***-0.01*** | ***0.02*** | ***18*** | ***-6780*** | ***1.9*** | ***0.3*** |
| 3 | -0.081 | 0.006 | | 0.001 | + | | + | | 0.02 | 0.00 | -0.01 | 0.02 | 24 | -6776 | 6.0 | 0.0 |
| 4 | -0.080 | 0.006 | 0.000 | 0.001 | + | | + | | 0.02 | 0.00 | -0.01 | 0.02 | 25 | -6774 | 8.1 | 0.0 |
| 5 | -0.081 | 0.004 | 0.002 | 0.001 | + | | | + | 0.02 | 0.00 | -0.01 | 0.02 | 25 | -6772 | 9.7 | 0.0 |
| 6 | -0.152 | 0.004 | | 0.001 | + | + | | | 0.02 | 0.00 | -0.01 | 0.02 | 24 | -6772 | 9.9 | 0.0 |
| 7 | -0.151 | 0.004 | 0.001 | 0.001 | + | + | | | 0.02 | 0.00 | -0.01 | 0.02 | 25 | -6770 | 11.9 | 0.0 |
| 8 | -0.062 | | 0.004 | 0.002 | + | | | | 0.02 | 0.00 | -0.01 | 0.02 | 17 | -6769 | 12.9 | 0.0 |
| 9 | | 0.003 | | 0.001 | + | | | | 0.02 | 0.00 | -0.01 | 0.02 | 16 | -6768 | 14.2 | 0.0 |
| 10 | | 0.003 | 0.001 | 0.001 | + | | | | 0.02 | 0.00 | -0.01 | 0.02 | 17 | -6766 | 15.6 | 0.0 |
| 11 | -0.062 | | | 0.003 | + | | | | 0.02 | 0.00 | -0.01 | 0.02 | 16 | -6766 | 15.6 | 0.0 |
| 12 | -0.170 | 0.007 | | 0.001 | + | + | + | | 0.02 | 0.00 | -0.01 | 0.02 | 31 | -6766 | 15.7 | 0.0 |
| 13 | -0.169 | 0.007 | 0.000 | 0.001 | + | + | + | | 0.02 | 0.00 | -0.01 | 0.02 | 32 | -6764 | 17.7 | 0.0 |
| 14 | -0.080 | 0.006 | 0.000 | 0.001 | + | | + | + | 0.02 | 0.00 | -0.01 | 0.02 | 32 | -6762 | 20.0 | 0.0 |
| 15 | -0.153 | 0.004 | 0.003 | 0.001 | + | + | | + | 0.02 | 0.00 | -0.01 | 0.02 | 32 | -6762 | 20.0 | 0.0 |
